# An RNAi screen reveals common host-virus gene signatures: implication for anti-viral drug discovery

**DOI:** 10.1101/2022.07.12.496853

**Authors:** David Shum, Bhavneet Bhinder, Jeni Mahida, Constantin Radu, Paul A. Calder, Hakim Djaballah

**Affiliations:** HTS Core Facility, Memorial Sloan Kettering Cancer Center, New York, USA

**Keywords:** HCS, HTS, genomic screen, shRNA, dengue virus, antiviral

## Abstract

Dengue is the most common mosquito-borne viral disease that in recent years has become a major international public health concern. Dengue is a tropical neglected disease with increasing global incidences, affecting millions of people worldwide, and without the availability of specific treatments to combat it. The identification of host-target genes essential for the virus life cycle, for which effective modulators may already exist, would provide an alternative path to a rapid drug development of the much needed anti-Dengue agents. For this purpose, we performed the first genome-wide RNAi screen, combining two high content readouts for DENV infection (FLUO) and host cell toxicity (NUCL), against an arrayed lentiviral based shRNA library covering 16,000 genes with a redundancy of at least 5 hairpins per gene. The screen identified 1,924 gene candidates in total; of which, 1,730 gene candidates abrogated Dengue infection, while 194 gene candidates were found to enhance its infectivity in HEK293 cells. A first pass clustering analysis of hits revealed a well orchestrated gene-network dependency on host cell homeostasis and physiology triggering distinct cellular pathways for infectivity, replication, trafficking and egress; a second analysis revealed a comprehensive gene signature of 331 genes common to hits identified in 28 published RNAi host-viral interactions screens. Taken together, our findings provide novel antiviral molecular targets with the potential for drug discovery and development.

## Introduction

The dengue viruses (DENV) constitute a single species within the genus *Flavivirus* of the *Flaviviridae* family including West Nile, and Yellow Fever virus; and transmitted to humans through infected mosquitoes. DENV infection begins with entry into cells by receptor-mediated endocytosis followed by replication of a single stranded, positive sense, RNA genome of approximately 10 kb in length. The RNA genome encodes for three structural proteins, involved in particle formation (capsid [C], membrane [prM], and envelope [E]), and seven nonstructural proteins (NS) involved in viral replication (NS1, NS2a, NS2b, NS3, NS4a, NS4b, and NS5)^3^ and all strains belong to one of four serotypes (DENV-1-4) which differ from each other by as much as 35% at the nucleotide level.^4^

According to the World Health Organization, DENV infection has emerged as one of the most significant health risks in tropical and subtropical regions worldwide in the past 30 years. It is estimated that upwards of 50 million people are infected yearly^1^ and two fifths of the world’s population is at risk for infection.^2^ DENV causes dengue fever and the more serious dengue hemorrhagic fever of which there are no vaccine or specific antiviral agent approved for use.

Previously, we describe the adaptation, validation and screening of our high content assay against a chemical library of known drugs and bioactives.^5^ Utilizing a multi-parametric readout to directly monitor viral replication in host cells, we simultaneously scored compounds for anti-viral activity that target several prominent classes of cellular factors including transporters, receptors, and enzymes. We are currently in follow-up studies to potentially repurpose these compounds for use as a new indication for antiviral therapies against DENV infections and perhaps other flaviviruses.

In an alternative strategy, we utilize RNAi technology to elucidate the mechanisms of viral entry and to understand how DENV recruits host proteins for each step of its life cycle. These critical factors remains largely unclear and their unraveling may eventually lead to the development of novel anti-viral therapies. RNAi technology has been used successfully to identify host protein involved in West Nile virus (WNV) infection.^6^ Kishnan et al identified 305 host proteins that affect WNV infection following a genome-wide siRNA screen in HeLa cells. Similarly, Konig et al performed a genome-wide siRNA screen and identified 295 co-factors required for influenza virus growth.^7^ Following on their success, we employ an arrayed approach using shRNA in the form of lentiviral particles to silence genes one hairpin at a time. An arrayed shRNA screen have several important advantages including stability of silencing rather than transient as with siRNA, efficient delivery, greater sensitivity to discern subtle phenotypes, and information-rich as those involving cell based assays with high-content imaging.^8^

In this article, we describe the adaptation of our assay from a chemical screening platform for screening against a genome-wide shRNA lentiviral particle library. Our assay utilizes a high-content multiparametric approach to evaluate the affect of host gene silencing on viral activity, assess cell cytotoxicity, and to monitor subtle phenotypes as a result of cellular changes. We report a comprehensive overlap with the already published results from 28 viral RNAi screens and the key gene, function and pathway signatures involved in host-viral interactions. Here, we present our methodology for screening, bioinformatics analysis, and hit characterization to gain a better understanding of host-viral processes in hopes developing new antiviral therapies against DENV and other flavivirus infections.

## Materials and Methods

### Cells, Virus and Antibodies

The cell line human embryonic kidney HEK293 (Microbix, Canada) was grown at 37°C in an atmosphere containing 5% CO_2_. HEK293 cells were propagated in complete growth medium containing Minimal Essential Eagle’s Medium (MEM) supplemented with 10% heat inactivated fetal bovine serum (FBS), L-glutamine, nonessential amino acids, 100 unit/ml penicillin, and 100 g/ml streptomycin. For assay development and screening studies, MEM was supplemented with only 10% heat inactivated fetal bovine serum (FBS), L-glutamine, and nonessential amino acids. All cell culture supplies were from Fisher Scientific and Invitrogen. DENV-2 strain New Guinea C was purchased from ATCC (American Type Culture Collection, Catalog #VR-222) and virus stocks were prepared as previously described.^5^

The DENV infection assay was industrialized for automated high throughput screening (HTS) in 384-well microtiter plate format as previously described. Polybrene tolerance, puromycin assessment, transduction studies, and assay development are described under Supplementary Methods.

### Genome-wide Arrayed shRNA Lentiviral Particle Screen of DENV Assay

HEK293 cells were seeded at 1,000 cells per well in 40 µL of growth medium into a 384-well microtiter plate and settled overnight. To prepare assay plates for transduction, 10 µL of growth medium containing polybrene was dispensed into the assay plate at a final concentration of 8 µg/ml. For transduction, shRNA lentiviral particles were pre-plated in 384-well polypropylene source plates containing ready-to-use shRNA lentiviral particles at 1×10^6^ TU per mL and 5 µL was transferred to the assay plates at a MOI of 5. Immediately after shRNA lentiviral particle addition, assay plates were centrifuged for 8 minutes at 1,300 rpm. After 96h incubation, transduction media was aspirated and 45 µL of growth media containing puromycin was dispensed to the assay plate at a final concentration of 1 µg/mL. After 168h incubation, selection media was aspirated and 35 µL of growth medium was added. DENV-2 virus were diluted in inoculation medium comprised of MEM supplemented with only 2% heat inactivated FBS, L-glutamine, and nonessential amino acids followed by 10 µL dispense to the assay plate at a MOI of 0.5. After 48h incubation, cells were fixed and immunostained for DENV E using the procedure as previously described.^5^ Images were acquired using the IN CELL Analyzer 3000 (INCA3000, GE Healthcare, USA) using a 40X magnifying objective allowing for nine images per well, covering 90% of the well and analyzed using Raven 1.0 software for DENV E and NUCL. The shRNA lentiviral particle library was plated in 295 384-well assay plates and required 1100h for imaging on the INCA3000.

### Image Acquisition and Analysis for the Genome-wide Screen

Image acquisition for screening was performed on the automated laser confocal INCA3000 microscope system. This laser scanning confocal imager comprises two laser light sources, three excitation lines and three highly sensitive 12-bit CCD cameras allowing simultaneous imaging of three fluorophores with continuous laser-based autofocus. Image acquisition was captured at the following wavelengths: 364nm excitation / 450nm emission in the blue channel for Hoechst stained nuclei and 488nm excitation / 535nm emission in the green channel for DENV E with an exposure time of 1.5 milli-seconds. Nine images per well collected using a 40X magnifying objective covering 90% of the well and required 32 seconds per well with total imaging time of 210 minutes for a complete 384-well microtiter plate. Images were analyzed using the Raven 1.0 software’s built-in object intensity analysis module. The object intensity analysis module uses a segmentation algorithm to determine NUCL with Hoechst pixel intensities above background. For DENV E, a cell masked overlay was generated using nuclei as a marker and intensity was measured within the defined boundaries. Analysis of NUCL and DENV E required 20 minutes for a complete 384-well microtiter plate.

### Statistical Analysis

Hit selection at the gene and shRNA level was performed using two methods: Control Based Method (CBM) and Ratio Based Method (RBM). First, The individual shRNA were selected as inhibitors of DENV infection if their DENV E intensities were less than mean + *2* standard deviation of the empty control wells. The genes were binned according to the number of active shRNAs and those with 3 or greater active shRNA were classified as hits. Second to evaluate cytotoxicity, individual shRNAs were scored as cytotoxic if NUCL count were comparable to puromycin control wells. Genes were filtered from the nominated hit list if 3 or greater shRNA scored as cytotoxic. For RBM, hits were nominated at the shRNA level using a ratio of DENV E intensities and NUCL. Individual shRNAs with ratios below our controls were scored as inhibitors. The genes were binned according to the number of scoring shRNAs and those with 3 or greater active shRNAs were classified as gene hits. Nominated hit lists from both CBM and RBM were combined into a gene list of DENV inhibitors. For the genes with less than 3 shRNAs in the library, the ones with a 100% hit rate were added to the hit list for inhibitors. (Hit rate is defined as the ratio between numbers of active shRNA to total shRNA for that gene). The genes were selected as enhancers of DENV E infection if their DENV intensities were greater than mean + *2* standard deviation of the wells with DENV alone. Statistical analysis for hit selection was performed using PERL scripts and Sigmaplot (SYSTAT, USA).

### Bioinformatics Analysis

The DENV inhibitor gene list was annotated with information from PANTHER (www.pantherdb.org/) and the Gene Ontology (www.geneontology.org). The gene lists were further manually curated with information obtained from Pubmed and UniProt (www.uniprot.org/). The canonical pathway associations and network building was performed in GeneGo’s Metacore pathway analysis software (www.genego.com/metacore.php). Metacore converts the gene ids into network objects for the analysis. The statistical level of significance used was p-value < 0.05 with a false discovery rate of 5%. The interactome functionally of Metacore was used to find overrepresented protein-protein interactions in the results. DAVID was used for functional clustering and the threshold p-value was set at < 0.05 and bonferroni correction. KEGGS, panther, reactome, biocarta and Wiki pathways were also used for pathway enrichments. The overlap analysis was performed using PERL scripts. The list of genes from the literature searches were unified by NCBI_IDs and gene synonyms obtained from Metacore.

## Results

### High-Content Assay to Identify Host Factors Involved in DENV Infection with shRNA

To identify host factors involved in DENV infection, our previously described high-content cell-based assay for compound screening was adapted for genomic screening based on shRNA technology in an arrayed format. Using our human model, HEK293 cells were transduced with shRNA lentiviral particles for 4 days followed by selection with puromycin for 7 days. Following gene silencing, HEK293 cells were infected with DENV-2 and 2 days later measured for amount of viral protein produced in the cells. A reduction in DENV E protein production would indicate that silencing of a particular gene or pathway potentially inhibits an important stage in the viral cycle including entry, replication, and processing. On the contrary, an increase in DENV E protein production indicates that gene silencing enhances viral production (**Fig. 1A**).

**Figure 1.**
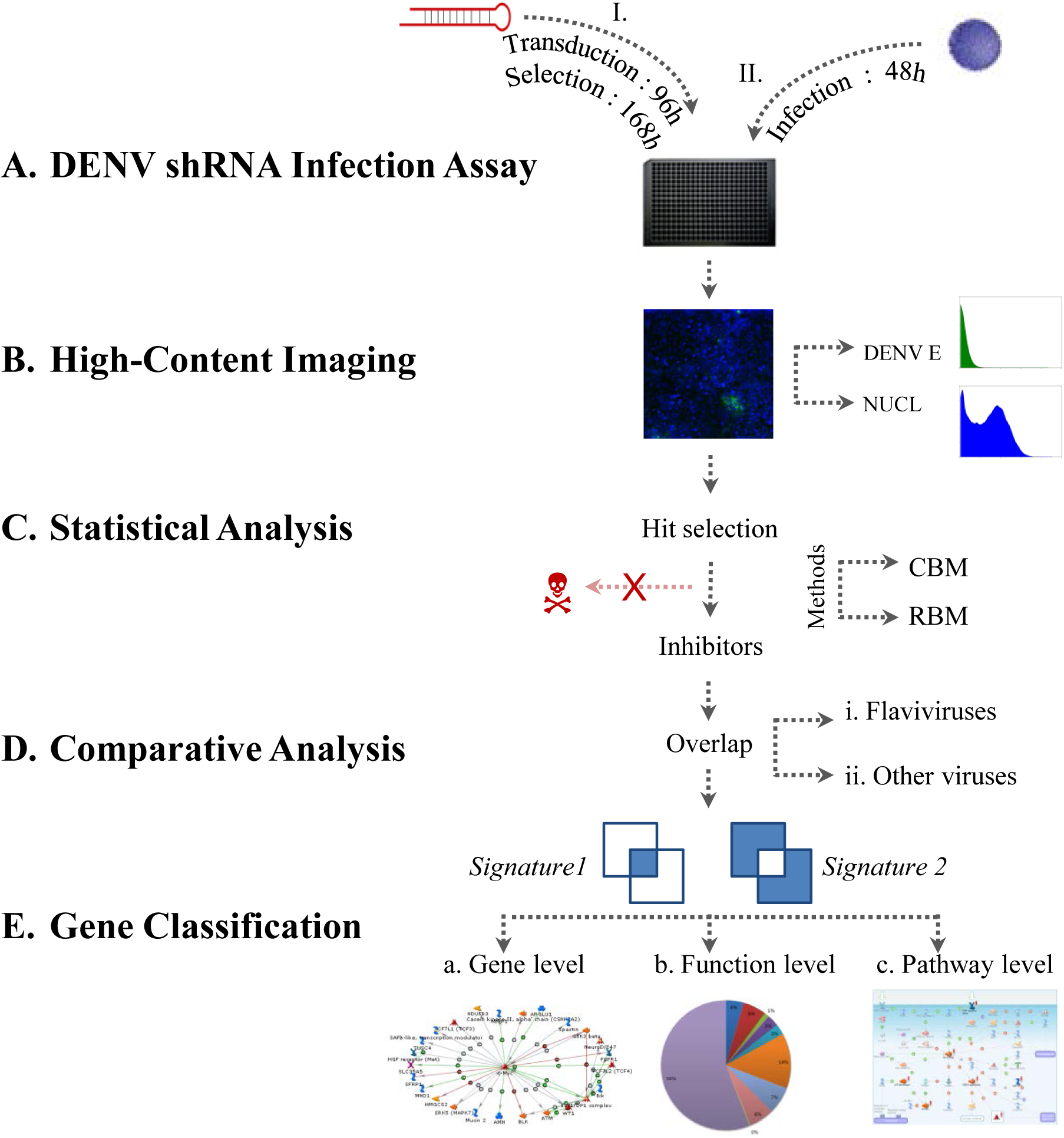
DENV shRNA infection assay and workflow.

By combining our high-content cell-based assay with shRNA arrayed approach, it allows for instantaneous evaluation of each gene at a single clone level without the need for deconvolution typical of many pooled RNAi screens. Using a multiparametric analysis, we are able to instantaneously evaluate gene silencing on viral activity, monitor host cell cellular changes, and assess cell cytotoxicity. For our genomic screen, we observed a DENV infection rate of only 10% in comparison with the chemical screen with DENV infection rate of over 90% of cells (**Fig. 1B**). The dramatic reduction in DENV infection is accountable with a first transduction event by lentivirus which disrupts the cell membrane and as a consequence; the second infection by DENV receptor mediated entry is affected. By utilizing a high-content based approach, we were able still able to extrapolate data by performing a cell by cell analysis for each shRNA. Similarly, Krishnan et al identified candidate genes by siRNA mediated silencing even though the WNV infection rate was between 20% - 30%.^6^

### Assay Development for shRNA Lentiviral Particle Screening

Using our previously described DENV infection assay for chemical screening as a guide, we focused on identifying the optimal conditions for genomic-based screening using shRNA lentiviral particles in a miniaturized 384-well microtiter plate format. We first investigated tolerance of HEK293 cells to polybrene, a cationic polymer used to enhance lentiviral transduction. Cells were seeded at densities between 500 and 2,000 cells per well in the presence of polybrene at 12-point doubling concentration series between 12 ng/mL and 25 μg/mL (**Supp. Fig. 1A**). Typically, lentiviral transduction process takes place over 4 days and we monitored growth by Hoechst stained nuclei imaging at 96h post seeding. At the polybrene concentrations tested, cells did not show slow recovery or cytotoxicity and 1,000 cells per well was selected as the initial seeding density with 8 μg/mL polybrene as the optimal concentration based on overall consistency of growth (**Supp. Fig. 1B**).

The shRNA lentiviral particle library used for screening is based on pLKO.1 vector system and contains a puromycin resistance marker for selection of positively transduced cells. We first investigated the sensitivity of HEK293 cells to puromycin in 12-point doubling concentration series between 12 ng/mL and 25 µg/mL. Afterwards, the IC_95_ was calculated as the puromycin concentration to effectively kill cells untransduced cells while selecting for positively transduced cells containing the resistance marker. To simulate the transduction process, cells were seeded at 1,000 cells per well with 8 μg/mL of polybrene for 96h followed by incubation with puromycin for 168h. Imaging of Hoechst stained nuclei was used to assess sensitivity of cells to puromycin and IC_95_ was determined by curve-fitting (**Supp. Fig. 1C**). We identified the IC_95_ of puromycin at 1 µg/mL as the selection criteria for positively transduced clones (**Supp. Fig. 1D**).

To evaluate the DENV assay for shRNA lentiviral particle screening, a panel of three control lentiviral particles was selected for testing: 1) a lentiviral particle containing scrambled shRNA was used to assess potential toxicity affects associated with transduction by interferon response; 2) a lentiviral particle containing TurboGFP was used as a visual monitor of transduction process; and 3) a shRNA lentiviral particle targeting PLK1^9,10^ an essential trigger for G2/M transition in which loss of expression results in pronounced aneuploidy was used as a phenotypic control to assess the transduction process. The control lentiviral particle containing scrambled shRNA was successfully transduced as depicted by consistency of growth of HEK293 cells. For the control lentiviral particle containing TurboGFP, we observed a dramatic up-regulation in green fluorescence indicating successfully transduction and expression of TurboGFP. For the control lentiviral particle containing PLK1, we observed enlarge cells with multi-nucleation as a phenotypic response consistent with aneuploidy as a result of gene silencing. In addition, we included a non-transduced well as a control and cells as expected showed pronounced cytotoxicity in the absence of resistant marker by puromycin selection (**Supp. Fig. 1E**).

### Screening of Genome-wide shRNA Lentiviral Particle Library

To identify host factors involved in DENV infection, the newly developed assay was as depicted in **Table 1** was screened against our genome-wide shRNA lentiviral particle collection in an arrayed format targeting 16,039 genes. The library was plated in 295 384-well microtiter plates with columns 13 and 14 empty for controls. To monitor the assay performance throughout the screen, control lentiviral particle containing scrambled shRNA was added to column 13 followed by DENV infection for high control and column 14 without DENV infection for low control (**Fig. 2A**).

**Figure 2.**
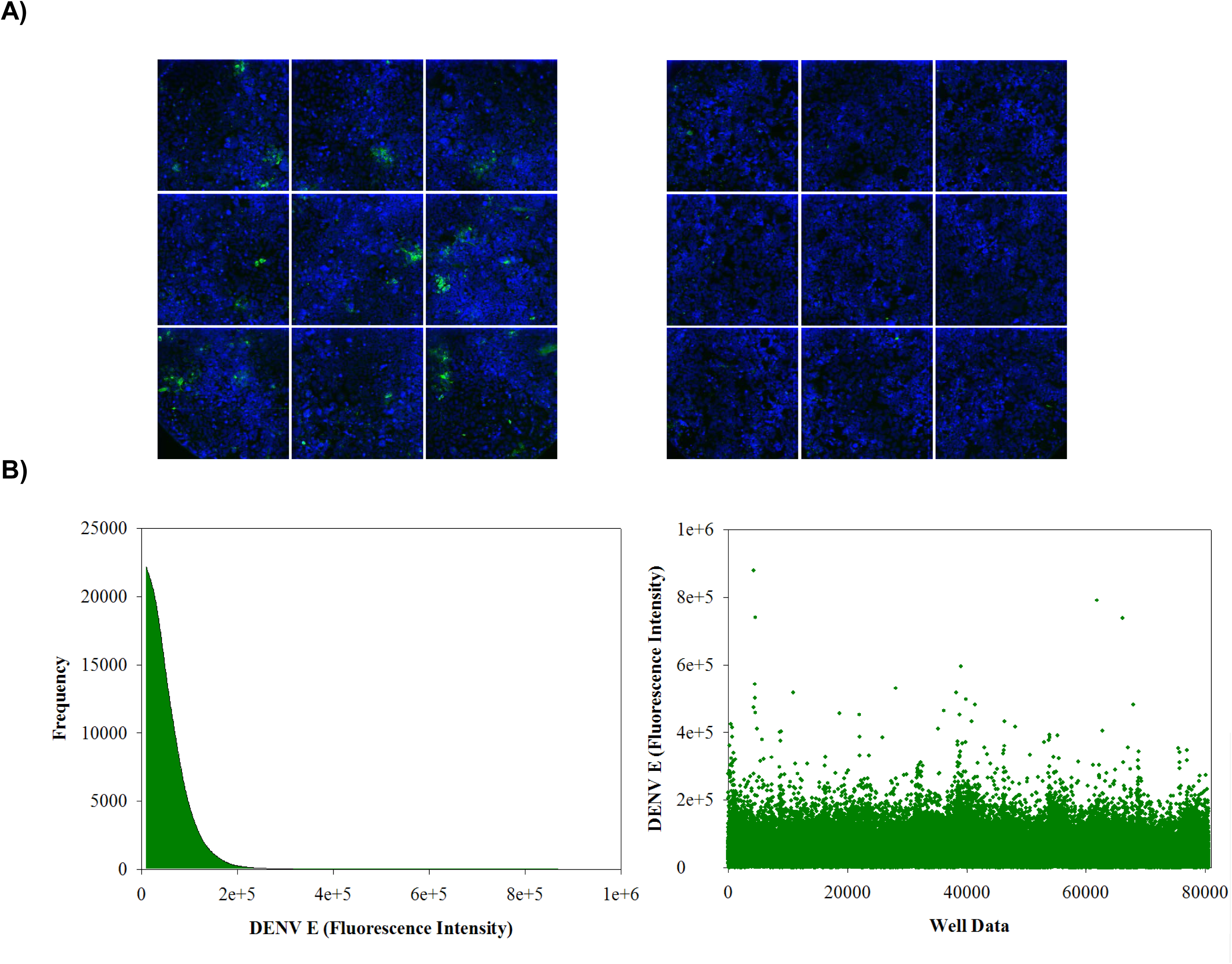
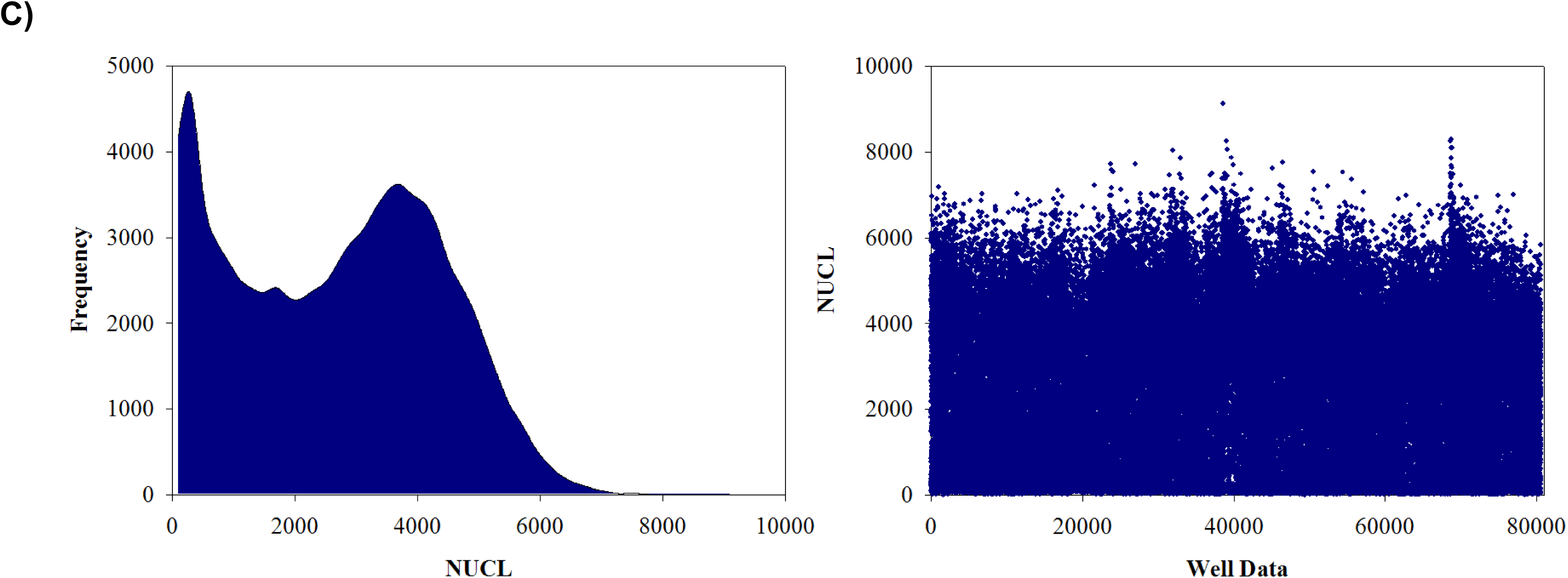
DENV shRNA infection assay screening results. **A)** Images were acquired using the INCA3000 with blue channel for detection of Hoechst stained nuclei and green channel for detection of DENV E. Left image: Sample high control well from column 13 of HEK293 cells infected with DENV. Right image: Sample low control well from column 14 of HEK293 cells without DENV. **B)** Frequency distribution of DENV E **C)** Frequency distribution of DENV E.

**Table 1.**
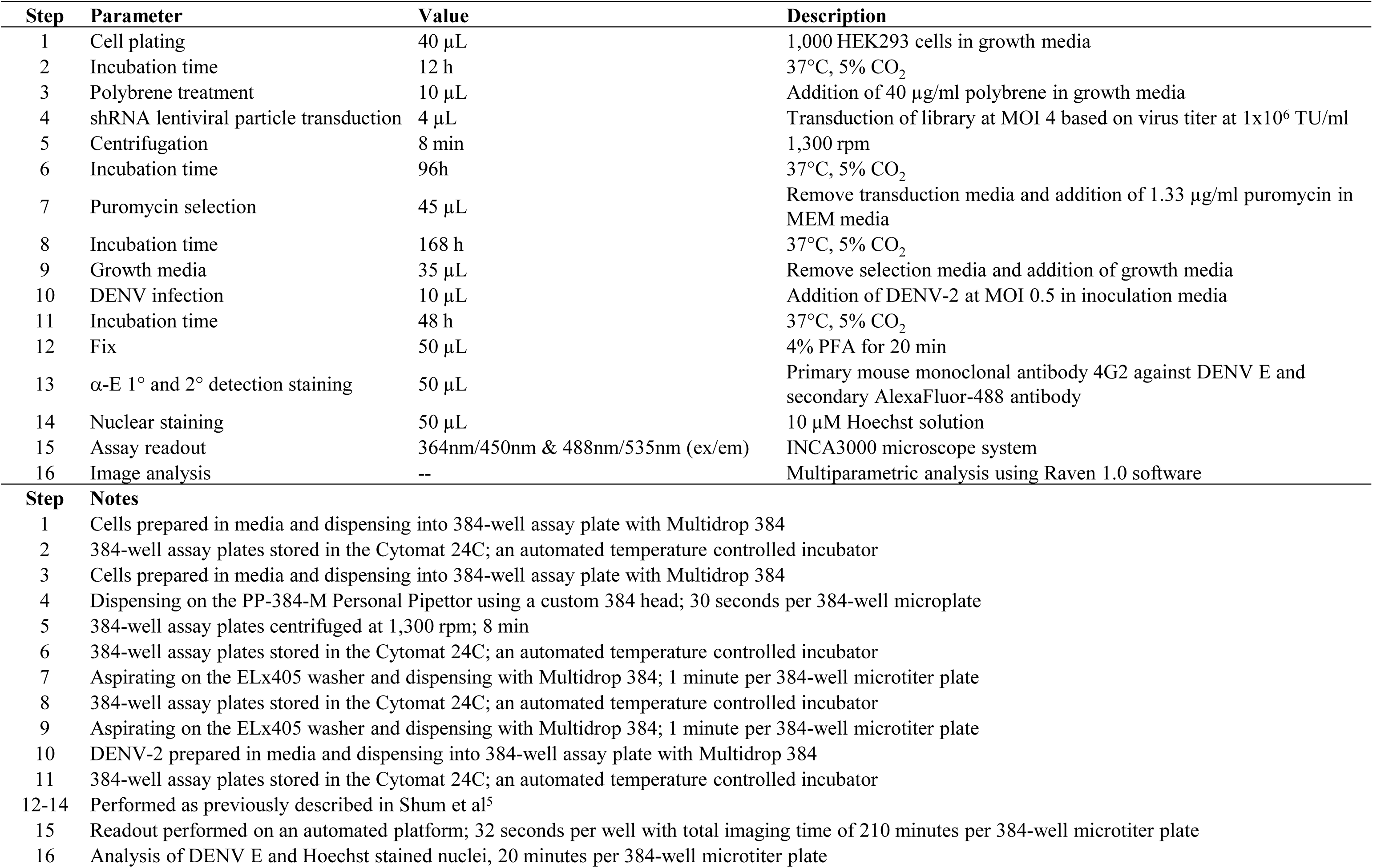
DENV shRNA infection assay workflow.

| Step | Parameter | Value | Description |
| --- | --- | --- | --- |
| 1 | Cell plating | 40 μL | 1,000 HEK293 cells in growth media |
| 2 | Incubation time | 12 h | 37°C, 5% CO<sub>2</sub> |
| 3 | Polybrene treatment | 10 μL | Addition of 40 μg/ml polybrene in growth media |
| 4 | shRNA lentiviral particle transduction | 4 μL | Transduction of library at MOI 4 based on virus titer at 1x10<sup>6</sup> TU/ml |
| 5 | Centrifugation | 8 min | 1,300 rpm |
| 6 | Incubation time | 96h | 37°C, 5% CO<sub>2</sub> |
| 7 | Puromycin selection | 45 μL | Remove transduction media and addition of 1.33 μg/ml puromycin in MEM media |
| 8 | Incubation time | 168 h | 37°C, 5% CO<sub>2</sub> |
| 9 | Growth media | 35 μL | Remove selection media and addition of growth media |
| 10 | DENV infection | 10 μL | Addition of DENV-2 at MOI 0.5 in inoculation media |
| 11 | Incubation time | 48 h | 37°C, 5% CO<sub>2</sub> |
| 12 | Fix | 50 μL | 4% PFA for 20 min |
| 13 | α-E 1° and 2° detection staining | 50 μL | Primary mouse monoclonal antibody 4G2 against DENV E and secondary AlexaFluor-488 antibody |
| 14 | Nuclear staining | 50 μL | 10 μM Hoechst solution |
| 15 | Assay readout | 364nm/450nm & 488nm/535nm (ex/em) | INCA3000 microscope system |
| 16 | Image analysis | -- | Multiparametric analysis using Raven 1.0 software |

We screened a library of 16,039 genes targeting 80,598 arrayed shRNAs for a dual readout of DENV E and NUCL. As anticipated, DENV E had an approximate normal distribution (**Fig. 2B**) while the NUCL followed a bimodal distribution (**Fig. 2B**), as typical to puromycin killing in shRNA screens. We used the CBM to score the reduction in DENV E intensity and found 11% of the total library genes with atleast 3 active shRNAs. For the second part of the analysis, we filtered out 313 inhibitors as cytotoxic. Next, we used the RBM to identify the non-cytotoxic inhibitors by analyzing each shRNA for DENV E intensity and cytotoxicity combined. These 2 methods together gave us an overall total of 1,730 inhibitor genes with a hit rate of 11%. Using the CBM to score the enhanced DENV E intensity, we identified 194 enhancer genes. Inhibitors were used in further analysis for signature identification.

### Inhibitors of Viral Infection from RNAi Screening Literature

To validate our screening results, we decided to study the overlap of our inhibitor list with the already published host viral interaction data. Since there are only ∼19 experimentally validated DENV-host gene interactions available^12^, we decided to expand our overlap analysis to include all viral RNAi screening studies published so far. To obtain this set of genes involved in host viral interaction, inhibitors from 28 other RNAi screening publications were obtained. These included 11 viruses - DENV, WNV, Human immunodeficiency virus (HIV), Hepatitis C virus (HCV), Influenza virus (IFV), Human papillomavirus (HPV), Epstein–Barr virus (EBV), Borna disease virus (BDV), Vesicular stomatitis virus (VSV), Marburg virus (MARV) and Sindbis. A summary of the studies selected and the differences in their methodologies are reported in (**Supp. Table 1**). A gene reported in at least 1 of the primary screen results was included in our analysis. During the comparative analysis we found that the only genome wide RNAi screen in DENV virus has been reported in Drosophila^11^. Thus, to our knowledge, our study is the only genome wide shRNA arrayed high content screening of DENV in human cells conducted so far. It’s noteworthy that there is a very poor overlap across the board and none of the host viral factors identified were consistently picked in all or most of the screens (data not shown). This observation is consistent with the overlap studies performed for the 3 HIV genome wide screens^13^. The genes thus obtained were further characterized into 2 groups; **set A** of 426 genes reported in *flaviviruses* (DENV and WNV) and **set B** of 2,920 genes reported in other viruses. For consistency, the gene names were standardized using their NCBI_IDs and gene synonyms obtained from Metacore. We noticed that 88 were common across all the viruses while 338 genes were unique to flaviviruses alone. Removing redundancies, 3,258 unique inhibitors were obtained which might be putatively involved in host viral interactions. An overview of the overlap across different studies is shown in **Fig. 3**.

**Figure 3.**
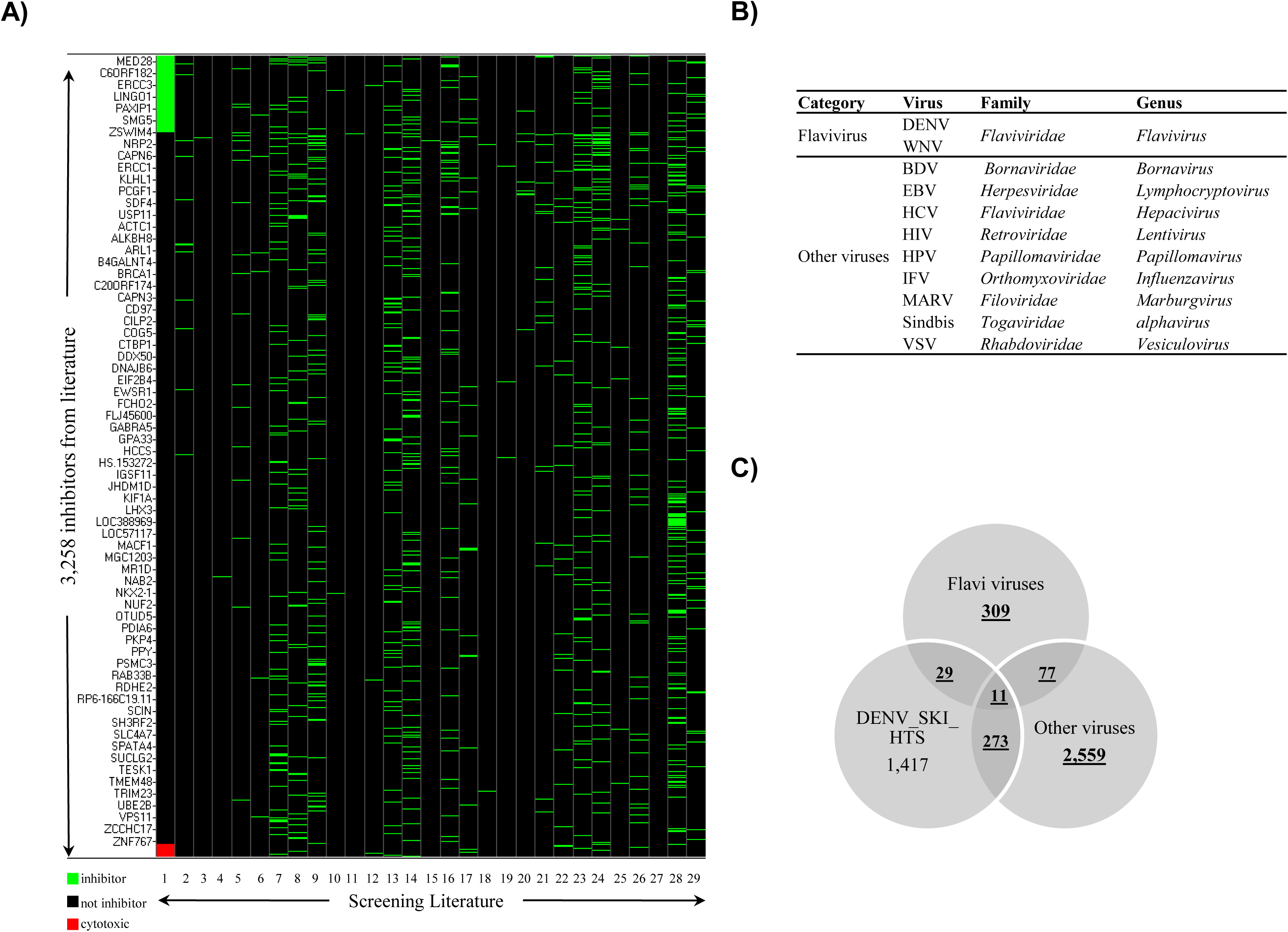
Overlap analysis across the 3,258 unique viral inhibitors reported in 28 RNAi screening studies for host-viral interactions. **A)** Heatmap showing the overall hit occurrence of the unique 3,258 inhibitors. Green represents active genes; black denotes inactive genes; red denotes factors tagged as cytotoxic in this study. The first column (1) of the heat map represents the hits identified in this study. For each of the following columns, the virus screened and the first author of the specific reference paper are : 2-DENV_Ang; 3-DENV_Heaton; 4-DENV_Rothwell; 5-DENV_Sessions; 6-DENV_Wang; 7-WNV_Krishnan; 8-HIV_Brass; 9-HIV_König; 10-HIV_Mao; 11-HIV_Nguyen; 12-HIV_Rato; 13-HIV_Yeung; 14-HIV_Zhou; 15-HCV_Berger; 16-HCV_Li; 17-HCV_Lupberger; 18-HCV_Ng; 19-HCV_Randall; 20-HCV_Suratanee; 21-HCV_Tai; 22-HCV_Vaillancourt; 23-IFV_Karlas; 24-IFV_König; 25-BDV_VSV_Clemente; 26-VSV_Panda; 27-EBV_MARV_Spurgers; 28-HPV_Smith; 29-Sindbis_Orvedahl. **B)** Classification of the different viruses used in the screening studies and their categorization into 2 groups which were further used in the analysis: flaviviruses and other viruses. **C)** Venn diagram representing the degree of overlap of inhibitors across 3 classes of hits namely flavivirus, other viruses and this study.

### Identification of Signatures Involved in Inhibition of DENV Infection

As reported in the previous section, using CBM and RBM, 11% of the genes from the screened library were selected as nominated hits. To validate the results of our screen, we did an overlap of our results with those obtained from literature in 2 parts- (a) inhibitors specific to flavivirus (set A of 426 genes) (b) inhibitors identified in other viruses (set B of 2,920 genes). We identified 40 genes from set A (**Supp. Table 2**) and 284 genes from set B to be common with inhibitors from our screen. An overlap between them revealed 11 overlapping inhibitors. These were primarily related to cytoskeleton, cell cycle regulation and immune response (**Table 2**). The genes which did not show any overlap with our screen were either inactive, cytotoxic, had < 3 active shRNA or absent in the library. A summary of the overlaps is shown in **Fig. 4**. Combining these results, we identified a total of 313 unique host-viral factors. It was interesting to note that 54 out of these were druggable targets (**Supp. Table 3**). Next, we analyzed the 313 common host-viral factors at 3 different levels – gene interactions level, functional level and pathway level. The following sections describe the results in detail.

**Figure 4.**
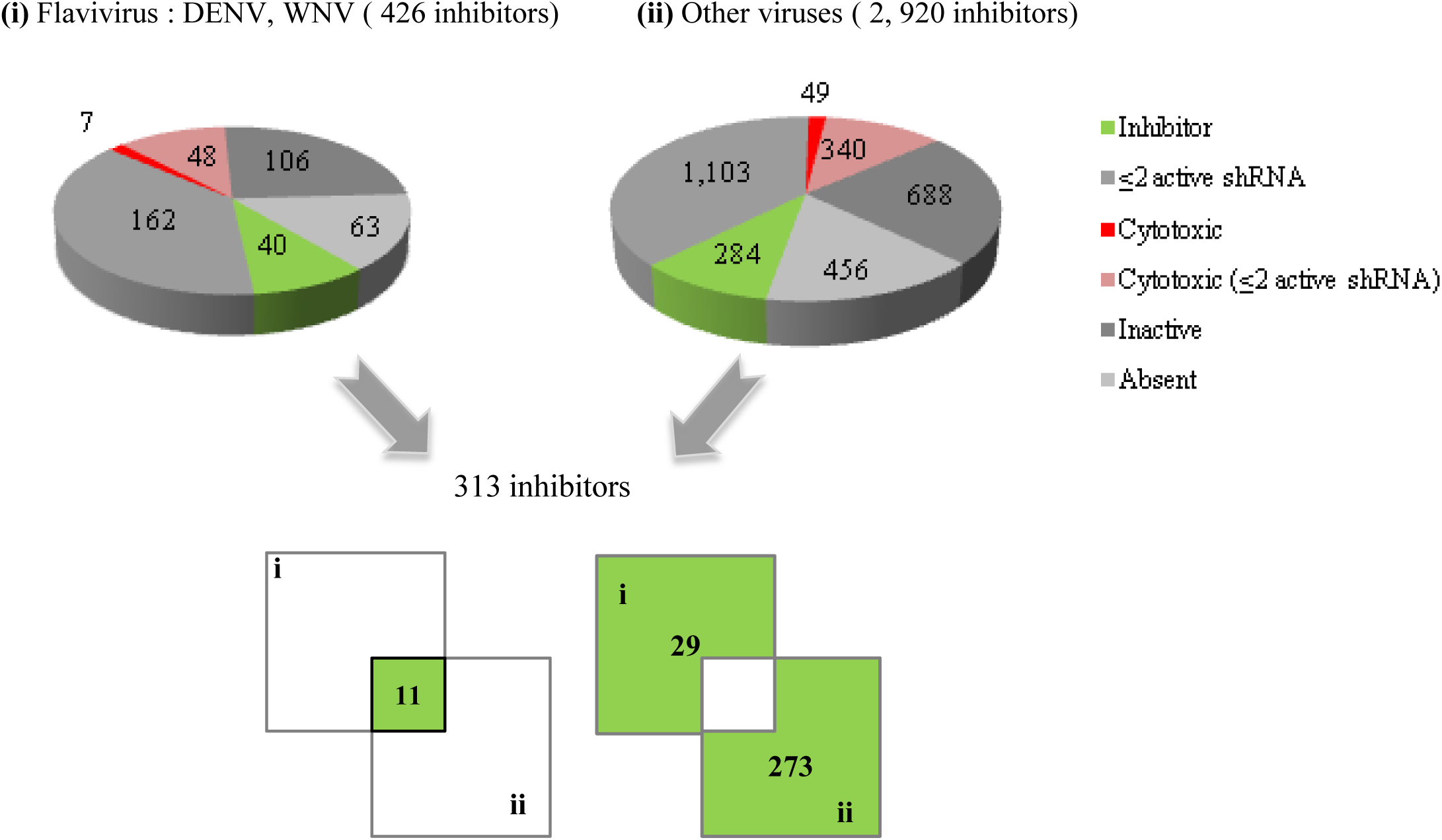
The workflow for selection of 313 unique core set of common inhibitors between this study and either of the 2 groups – flavivirus and other viruses. The hits overlapping between this study and each of the virus group are represented in green while the cytotoxic genes are represented in red. A breakup of the remaining genes not selected is shown in grey.

**Table 2.** The 11 genes common across 3 three groups – flaviviruses, other viruses and this study

| Gene Symbol | Description | Biological function | Family | Localization |
| --- | --- | --- | --- | --- |
| CALCB | calcitonin-related polypeptide beta | Vasodilatation inducer, calcium homeostasis | Calcitonin | Secreted |
| CFHR5 | complement factor H-related | Complement activation | Complement factor H | Secreted, extracellular |
| CSNK2A1 | casein kinase 2, alpha 1 polypeptid | Phosphorylation,, nucleo-cytoplasmic shuttling of transcription factors, DNA repair, Wnt signaling | Ser/Thr | Cytosol |
| GLCCI1 | glucocorticoid induced transcript 1 | Uncharacterized; interacts with VPS28 ( endosomal sorting) | ? | Nucleus, Cytoplasm |
| KIF17 | kinesin family member 17 | Vesicular transport | Kinesin | Microtubules |
| PLK4 | polo-like kinase 4 | Regulates centriole duplication during the cell cycle( G2/M transition) | Ser/Thr | Centrioles |
| PTK2B (FAK2) | protein tyrosine kinase 2 beta | Calcium-induced regulation of ion channels, MAPK signaling | Tyr | Cytoskeleton, focal adhesion |
| ROCK1 | Rho-associated, coiled-coil containing protein kinase 1 | Regulates formation of focal adhesions, adhesion & aggregation of platelets, essential in cytokinesis, transcriptional activation, stabilization, apoptosis, signal transduction | Ser/Thr | Cytoskeleton, Golgi |
| ROCK2 | Rho-associated, coiled-coil containing protein kinase 2 | Regulates cytokinesis, formation of actin stress fibers & focal adhesions, activation of the c-fos serum response element | Ser/Thr | Cytosol, microtubule |
| TNFSF13B | tumor necrosis factor (ligand) superfamily, member 13b | B-cell activator, immune response, B-cell proliferation, signal transduction | TNF | Membrane, secreted |
| WDR18 | WD repeat domain 18 | Cell cycle progression, signal transduction, apoptosis, gene regulation | WD repeat | Nucleus(?) |

### a) Gene Hubs as the Key Gene Signatures

The interactome functionality of metacore was used to identify the statistically significant gene-gene interactions within the dataset. The over-connected genes (hubs) were characterized as signature genes, which could be exploited as potential drug targets to combat viral infections. We identified 333 significant gene-gene interactions within the dataset, and found 31 gene hubs with 4 or more direct protein-protein (pp) interactions. Amongst the ones with maximum interactions were 11 hub genes: MYC (44 pp interactions), EGFR (24 pp interactions), p300 (29 pp interactions), E2F1 (18 pp interactions each), and AKT1, EZH2, TRIM28 (14 pp interactions each), CDK2 (13 pp interactions), CDKN1B/p27kip1 (11 pp interactions) and APC, GSK3B (10 pp interactions each) (**Fig. 5A**). Notably, 10 of these genes are predominantly regulators of various stages of cell cycle regulation while TRIM28 is involved in transcription regulation. The number of active shRNAs in our results for these genes are; EGFR (7 shRNA), EZH1 (5 shRNA), MYC (5 shRNA), p300 (5 shRNA), CDKN1B (4 shRNA), E2F1 (4 shRNA), TRIM28 (4 shRNA), AKT1 (3 shRNA), APC (3 shRNA), CDK2 (3 shRNA) and GSK3B (3 shRNA). Next, we studied the interaction of the hub genes with the experimentally validated DENV-host genes through the network functionality of Metacore. We found that 8 hub genes (AKT1, CDK2, CDKN1B, E2F1, EGFR, MYC, p300 and TRIM8) had a direct and 3 had indirect interactions in a network with 16 DENV-host genes (**Fig. 5B**).

**Figure 5.**
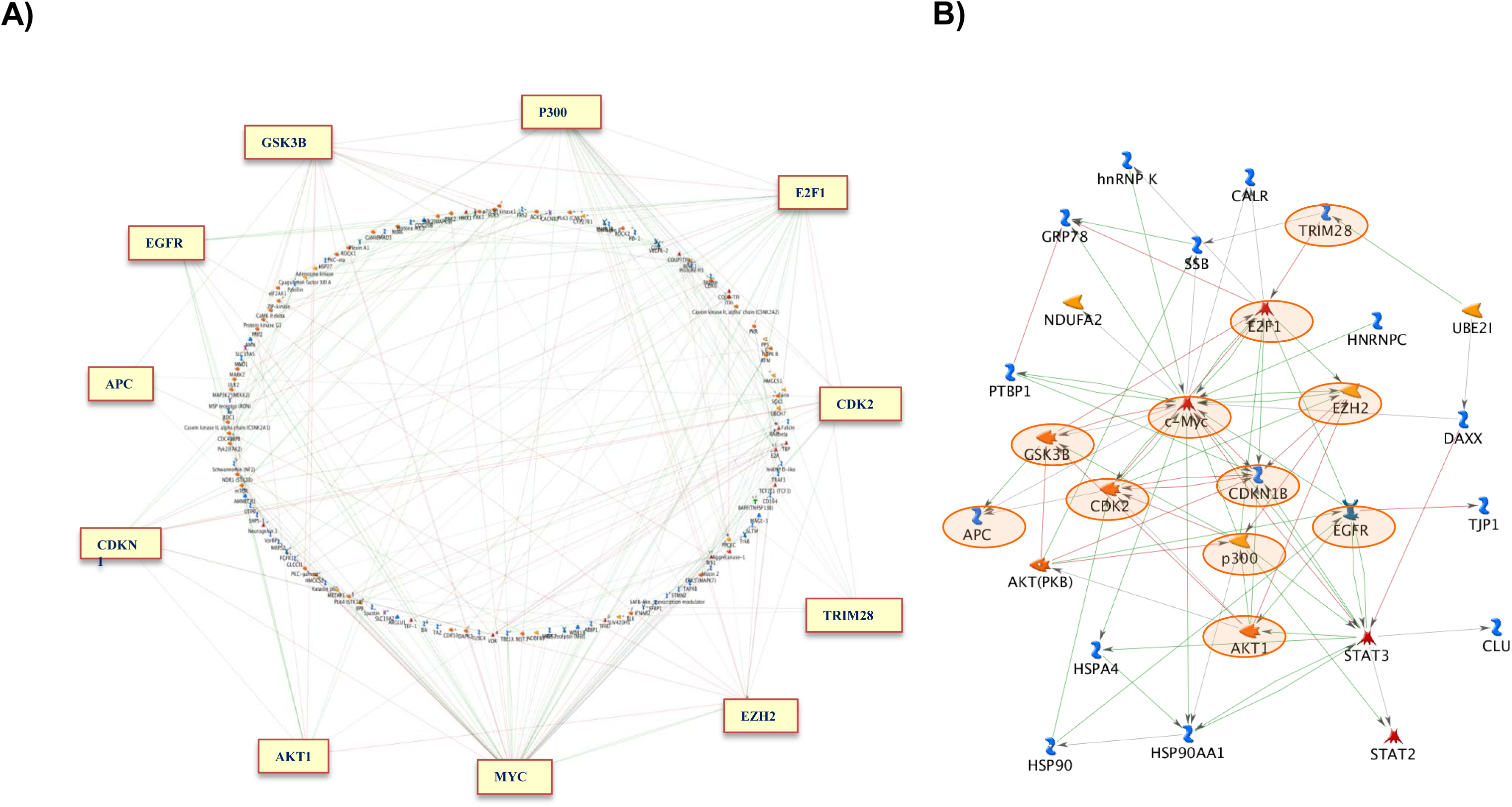
Gene signatures as over connected genes in the dataset. This is based on the interactome result of the 313 inhibitors obtained from Metacore. **A)** The 11 most densely connected genes (>10 protein-protein interactions) are represented in this graph. **B)** The interaction of the hub genes with the experimentally validated targets of dengue infection. The hub genes are highlighted with circles in the figure.

Furthermore, we explored the direct or indirect role of the hub genes implicated in other viruses and found that 8 to be involved in other host-viral interactions. It has been previously shown that MYC expression and stability is associated with HCV infection^14,15^ . Also, translation factor eIF4E, modulated in DENV infection^16^, is under the positive regulation of MYC^17^. The signaling mediated by EGFR is involved in protein phosphorylation and subsequently in control of cell proliferation and differentiation. EGFR has been reported to be involved in internalization of HCV via membrane fusion ^18^. The activation of EGFR for viral entry has also been experimentally validated in cytomegalovirus^19^. This observation suggests a putative role of EGFR in DENV internalization, which enters host cell via clathrin, mediated endocytosis, a mechanism similar to HCV entry^18^ (. p300 is involved in cell cycle, DNA damage and interferon immune response. Although not reported in flaviviruses, it has been shown that p300 mediated acetylation activates viral transcription in HIV-1^20^ and HPV^21^. Activation of transcription factor E2F1 is involved in HIV neuropathogenesis^22^ and infection of BHV-1 and HSV-1. It has also been shown that E2F1 and CDK2 get activated in HCV^23^ and HIV^24^. AKT1, an important anti apoptotic factor, might have a role in promoting the latent stage of the viral life cycle and studies which inhibit Akt report reactivation of viral lytic cycle in Herpes virus^25^. Akt signaling pathways have been shown to activated during infections by Influenza^26^ and Epstein-bar virus^27^. Also, another hub gene, GSK3B, is under the regulation of Akt, and acts in transcription regulation via the Wnt-3/beta-catenin signaling pathway. GSK3B is also associated with endoplasmic reticulum (ER) Stress overload pathway. CDKN1B is proposed to have a role in HCV^28^. To our knowledge, the remaining 3 genes (EZH2, TRIM8, and APC) have not been proposed to have a role in viral infection progression. However, it was interesting to note that TRIM8 is indirectly involved in activation NF-κB pathways^29^, a pathway modulated in the early stages of viral infections.

Also, the data obtained from drug bank revealed that AKT1, CDK2, EGFR and GSK3B have either a small molecule or biotech drugs available, thus highlighting their potential in antiviral drug development.

### b) Gene Classes as the Key Function Signatures

We explored various GO processes that were associated with our genes of interest. Using the PANTHER annotations, we were able to broadly classify our hits into molecular functions (**Supp. Fig. 2A**) and biological functions (**Supp. Fig. 2B**). To explore the biological functions in more detail, we performed the functional clustering in DAVID. We were able to identify enriched GO categories (p-value < 0.05) within the data set including signal transduction, cell cycle regulation, cytoskeleton organization, MAPK signaling and apoptosis (**Fig. 6A**). However, due to the hierarchy of GO terms, we also obtained a class of broad categories like metabolic process, protein phosphorylation, and cellular and biosynthetic process. To address this issue of innate redundancy, we utilized the gene lists manually annotated with information from NCBI and UniProt databases. Thus, by combining information from GO terms and manual curation, we were further able to refine the enriched biological processes into 18 functional signatures (**Fig. 6B**) and list of the genes in each signature are shown in **Supp. Table 4.**

**Figure 6.**
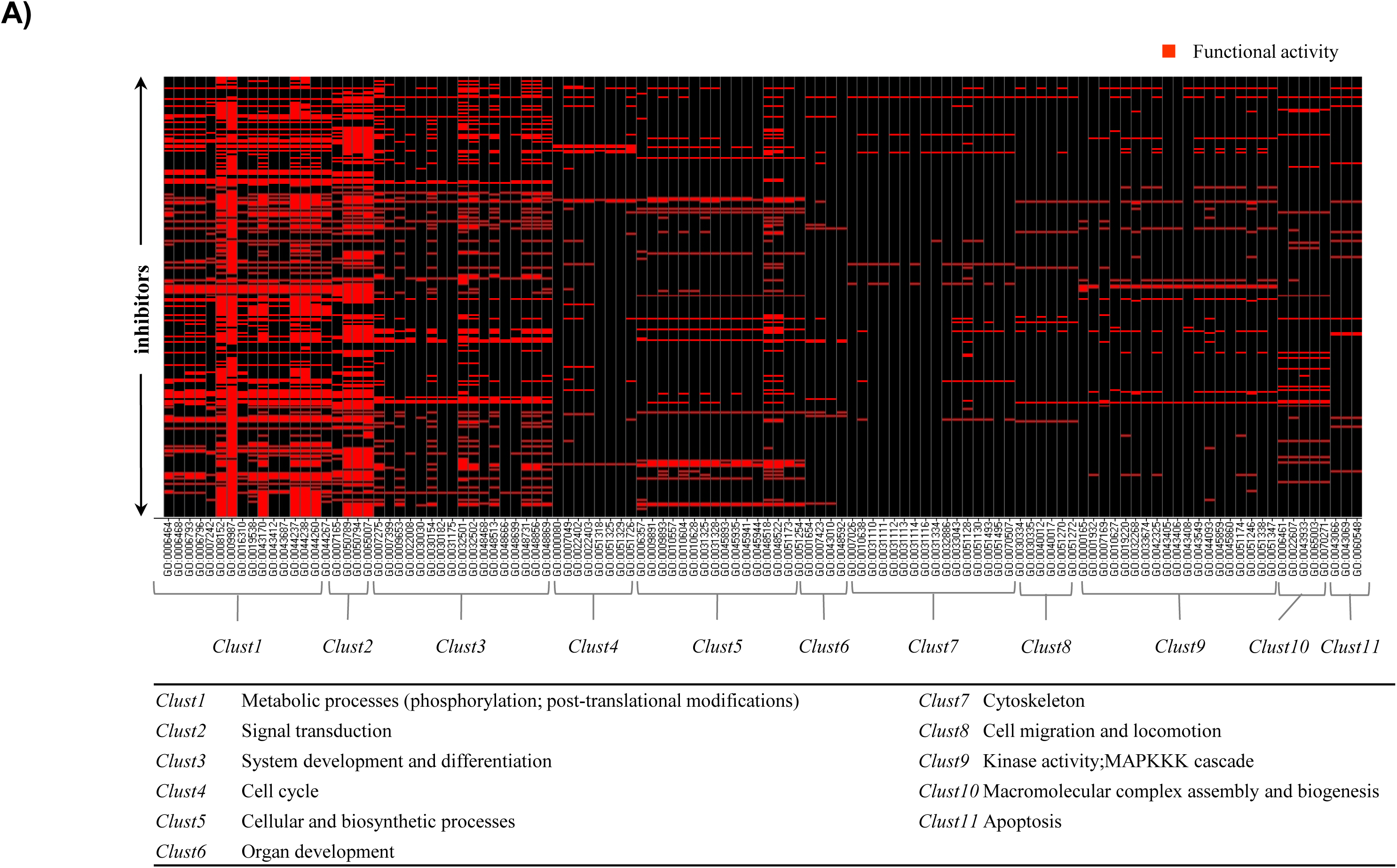
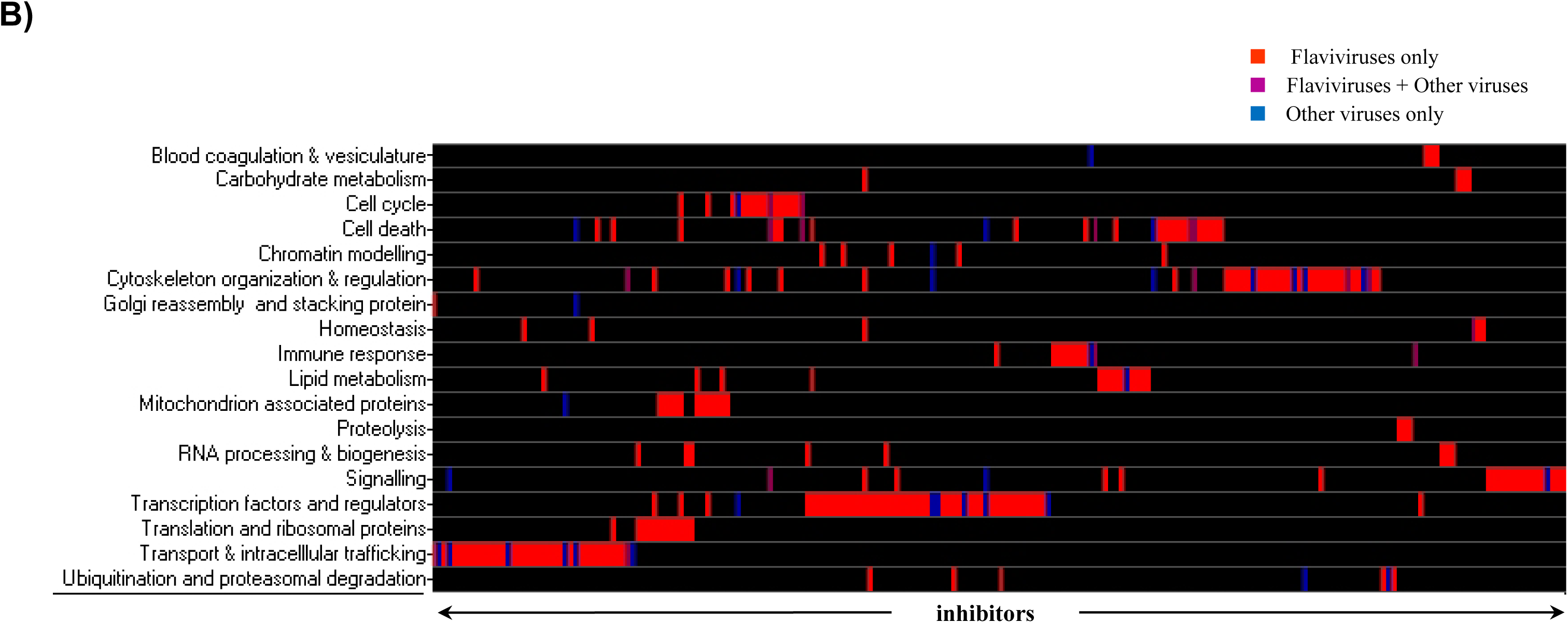
Functional profiling of 313 common inhibitors. **A)** Functional clustering of the enriched GO terms for the 313 common inhibitors into 11 clusters obtained using DAVID. Each cluster contains GO terms with a p-value < 0.05. The major function associated with each cluster is shown in the table beneath. Red denoted the genes active in the corresponding GO categories. **B)** 18 functional signatures found in the 313 genes. This graph is based on the gene annotations obtained from NCBI and UniProt databases. The black denotes functional inactivity of the corresponding genes.

We identified a total of 51 genes, which were either transcription factors or regulators including 2 members of the TFIID complex (TBP, TAF4B, TBPL1) and hnRNPs (HNRNPAB, HNRPDL). 13% of the genes were involved in cytoskeleton reorganization and regulation. The cytoskeleton reorganization genes involved actin, microtubule and myosin genes. The third most enriched category with 12% of the genes was that of transport and intracellular trafficking. This category included genes involved in endocytosis (RAB5B), exocytosis (RAB3D) and retrograde transport (COPG, DCLK1). We identified 10 ion transporters including calcium (CACNG4, CACNB2, STIM2), potassium (KCNAB2, KCNK4) sodium and magnesium ion transporters. This observation is consistent with our previous chemical screen in DENV (5). Besides, we also identified transporters for macromolecules like proteins, lipids and sugars.

Furthermore, we identified key genes associated with mitochondria (NDUFS3) and Golgi apparatus (GORASP1, GORASP2). Another set of structural protein identified was the 5 ribosomal proteins (MRPS2, MRPS7, MRPS10, MRPS12 associated with mitochondria and RPS6KB1). A major group of genes was found related to cell cycle regulation especially for progression of G1 phase, chromatin modeling, translation regulation, RNA processing, mRNA transport (IPPK), ubiquitination and proteosomal degradation. Another set of signatures was associated with the cell stress response including immune responses (complement system) and cell death including programmed cell death (PDCD1), autophagic (ULK2) and apoptotic genes. As another functional category, we identified 5 genes (VEGFR, FI3A1, H2F3A, SIRPA, TYMP) associated with blood coagulation, angiogenesis and vasculature. The functional signatures thus obtained can map the use of host cellular machinery during all stages of the viral life cycle inside the host cells.

### c) Canonical Pathways as the Key Pathway Signatures

The pathway signatures could be mapped to the various stages of the viral life cycle from entry to egress. The enriched pathways identified could be classified into 7 broad categories – cell cycle regulation, transcription, translation, cytoskeleton, clathrin-mediated endocytosis, lipid metabolism and signaling pathways.

The Metacore pathway analysis revealed the top 10 scoring pathways and their respective p-values as – cell cycle regulation: G1/S transition (2.5e-^10^), cytoskeleton remodeling (8.7e-^08^), transcription: p53 signaling pathway (1.5e^-06^), translation: regulation of EIF2 activity (1.5e^-06^), signal transduction: PI3K/AKT signaling (3e^-06^); VEGF signaling (4.2e-^06^); PTEN pathway (4.8e^-06^); PIP3 signaling (5.6e^-06^); EGFR signaling (4e^-05^) and cell adhesion: chemokines (1.7e^-05^). Next we explored other pathway resources available through DAVID (Keggs, Panther, Reactome, Biocarta) and found few additional enriched pathways which include signaling pathways; ErbB (2e-05), MAPK (2.62e^-04^); Wnt (5.1e^-04^), Phosphatidylinositol signaling (8.4e-2), Cytoskeleton: focal adhesions (3.5e-5); axon guidance (2.5e-2).

We found 5 genes (CDK2, CDK6, E2F1, GSK3B, ATM, KIP1) involved in cell cycle regulation mainly associated with cell cycle checkpoints and DNA repair. Also, it has been shown that flavivirus increases expression of TAP1 which is under transcription regulation of p53^30^. The 2 key members of the EIF2 pathway (EIF2AK1, PKR) were identified that phophorylate and inhibit EIF2 mediated translation. PKR is specifically activated by interferon in stress e.g., viral infection^31^

Furthermore, we picked up multiple pathways and the receptors, which initiate these pathways (EGFR, FGFR, VEGFR, MET). PI3K/AKT pathway is an anti-apoptotic pathway with a putative role in DENV^32^ . A major player of this pathway, AKT, also identified as a signature gene, is crucial mediators of BAD mediated apoptosis. It also regulates GSK3, a protein kinase, involved in glycogen synthesis and transcription regulation. Furthermore, we identified 16 genes in the MAPK signaling canonical pathway (CACN, TRKA/B, EGFR, FGFR, PKC, Mos, cPLA2, MNK1/2, c-myc, JNK, AKT, HSP27, PP5, PAK1/2, MEKK2B and ERK5), 11 genes in the ERBB pathway (Erb1, PKC, FAK, PAK, JNK, Myc, RAF, p70S6, GSK3B, AKT, KIP1), and 5 genes in the phosphatidylinositol signaling (PI4K, PIPK, PKC, IPK, DGKB). Another pathway of interest is the Wnt canonical signaling where we identified key members of the scaffold (CK2, GSK3B, APC, CK1 alpha), FRP, CREBBP, TCF4 and Lef. Also, we found P53 signaling pathway (APC and Ebil) involved in Proteolysis. It can be noticed that some genes were redundant in multiple signaling pathways, suggesting the interplay within these pathways to regulate apoptosis, cell cycle, transcription and translation. VEGF pathway is involved in vascular permeability, angiogenesis and cytoskeleton remodeling. Also, it is noteworthy that most of these pathways are under the influence of calcium ion concentrations in the cell. In the cytoskeleton remodeling category, we found genes in actin and microtubule organization and regulation e.g., COL4A6 (part of complex of the major structural component of basement membranes), PXN (cytoskeleton protein), TUBAL3 (microtubule component) ROCK1 and ROCK2 (regulators of actin organization). Although we did not significant enrichment in pathways related to immune response, we did find p300, VDR, and CREBBP^33,34^, which act in modulation of immune response.

Since the available tools rank the pathways by p-values, it is possible that some biologically relevant pathways, which fell below the threshold, might be missed. To combat that limitation, based on functional signatures and manual curation, we found genes associated with clathrin-mediated endocytosis (6 genes) and lipid metabolism. Thus, combining these observations, we identified signature pathways that could be modulated during viral infection. **Supp. Fig. 3** exemplifies some of the representative inhibitors that are members of some of these key canonical pathways for potential consideration for antiviral drug discovery.

## Discussion

DENV, an arthropod borne virus with a small 10.2k genome and encodes only 10 proteins, 3 structural and 7 non-structural proteins. Intuitively it requires multitude of host factors during its life cycle. Inside a host, the first stage in the infection cycle is receptor-mediated endocytosis of the virions. After viral ingestion, the virions undergo repeated cycles of replication and translation in the ER. The immature virions form a lipid coat and undergo protein folding and maturation in the trans-golgi network. After the latter process of virion assembly, the viral egress begins by modulating the permeability of the plasma membrane^35^. The entry of the virus in the host cell triggers a cascade of cellular processes at all stages of its progression including stress responses, membrane trafficking and signaling, cytoskeleton remodeling and vesicular transport. Although the DENV life cycle is categorized yet little is known about the complex host-viral interaction. Since the knowledge of host-DENV interactions is very limited, prediction-based studies for genome wide host-viral pp interactions have been reported^12^. However, such predictions still need to be experimentally studied and validated.

In our approach, we conducted the first genome-wide shRNA screen to identify the modulators of DENV infection and identified 1,730 inhibitors and 194 enhancers. We conducted an overlap with 28 other screening studies and found 313 common inhibitors, 40 of which overlapped with flaviviruses; out of which 11 were also overlapped with other viruses. This analysis helped us validate our results and introduce key signatures that are contained within the host-viral domain, thus giving us high confidence genes for further investigation. The signatures resonate the pathways and processes proposed to be involved in host-viral interactions by various published studies.

Microtubules and actin filaments have been previously shown to be actively involved in viral internalization, transport and egress and these movements might be regulated by signaling pathways e.g., MAPK, ERK^36,37^. Consistent with those findings, we found signatures enriched in cytoskeleton remodeling and related signaling pathways. It has been previously reported that flaviviruses enter host cells via clathrin mediated endocytosis^38^. We identified some key members of this pathway including Rab proteins, further supporting this phenomenon. Moreover, since this is a receptor mediated process and based on the findings in other viruses, we hypothesis a key role of EGFR in viral entry via endocytosis^18,19^. In addition, we also identify a Na+/H+ exchanger SLC9A6 with a putative role in acidification of endocytic vesicles^39^.

In an observation consistent with our previous chemical screen, we found calcium ion channels (CACNG4, CACNG4, and TRPV5). These, along with actin dynamics, might be involved in calcium-regulated exocytosis of mature virions. Also, the potential involvement of calcium ion influx and RAB3D in regulation of DENV exocytosis can be further investigated. We also identified STIM2, a likely candidate of Store-Operated Channels^40^ that regulates the inflow of calcium into the cell on ER reserve depletion. Also, it is noteworthy that the calcium ions in the cell regulate various signaling process including transcription regulation via Calcium-clamoudin pathway. Hence, it can be proposed that calcium ion channel might have a putative role in DENV infection and should be further explored as therapeutics.

The role of mitochondrial functions in DENV virus infections has been studied^41^. On the same lines, we identified the member of the first enzyme complex in the electron transport chain, NDUFS3 which maintains the cellular ATP levels through ATP synthase. Furthermore, we identified an ATPase inhibitor, ATPIF1, which inhibits ATP hydrolysis during depletion of the electrochemical gradient^42^. We also identified mitochondrial ribosomal proteins, although the function of these proteins is not well known, but they might be involved in oxidative phosphorylation and apoptosis^43^.

Although the role of cell cycle in DENV is not well known, yet we were able to find an enrichment of cell cycle genes in our analysis. Sine these genes are mainly connected to the G1/S phases and given that cell is under stress due to infection, it can be hypothesized that these genes regulate steps of DENV translation initiation via the internal ribosome entry site (IRES) mechanism^44^. We also identified host genes involved in viral transcription and protein folding and maturation.

Fatty acid biosynthesis and redistribution affected during DENV infection and might have a role in ER membrane expansions or coating of immature virions^45,46^. We have identified multiple factors involved in lipid metabolism, transport and homeostasis. Interestingly, we have identified HMGCS1, a rate limiting enzyme of the cholesterol biosynthetic pathway, which catalyzes conversion of acetyl-CoA to HMG-CoA. It has been previously shown that hymeglucin, a small molecule inhibitor of this enzyme reduces DENV infection, thus further supporting our finding^46^.

In agreement with earlier studies, we identified sets of genes associated with stress response, which can be characterized into cytopathic responses including anti- and pro-apoptotic factors and autophagy. Autophagy has emerged with a role in DENV replication via regulation of lipid metabolism^47^. We have identified ULK2 and autophagy regulator mTOR, further supporting this finding.

In conclusion, we present a comprehensive overlap with other RNAi screening studies and identified key gene, function and pathway signatures representative of the host-viral interactions with a focus on DENV. These signatures provide a high confidence set of DENV highlight key process and inhibitors affected during viral infection, 54 of which are druggable targets, and can further be investigated for experimental validation, and subsequently, antiviral drug discovery and development. However, there is an urgent need for novel anti-dengue drugs with the hope that our findings could facilitate the discovery.

## Acknowledgements

The authors wish to thank members of the HTS Core Facility for their help during the course of this study.

## List of Abbreviations

DENV: Dengue virus
shRNA: short hairpin RNA
DENV E: fluorescence intensity of Dengue virus glycoprotein envelope
NUCL: nuclei count
INCA2000: IN CELL Analyzer 2000
INCA3000: IN CELL Analyzer 3000
WNV: West Nile Virus
HIV: Human immunodeficiency virus
HCV: Hepatitis C virus
IFV: Influenza
BDV: Borna Disease Virus
VSV: Vesicular stomatitis virus
EBV: Epstein-Barr Virus
MARV: Marburg virus
HPV: Human papillomavirus

## Supplementary material

### Supplementary methods

#### Liquid Dispensing and Automation

Several different liquid dispensing devices were used in this study. The shRNA Lentiviral Particle Library was transferred from source plates (ABgene, UK) to assay plates (Corning, USA) using a custom-designed 384 head on a PP-384-M Personal Pipettor (Apricot Designs, USA). The addition of cell and virus suspensions was performed using the Multidrop 384 (Thermo, Canada) and assay plates were incubated in the Cytomat 24C (Thermo, Canada) under controlled humidity and an atmosphere of 5% CO_2_ at 37°C. Cell fixation and immunostaining were performed using the Biotek ELx405 washer (Biotek, USA).

#### Polybrene Tolerance Studies for shRNA Lentiviral Particle Assay Development

For polybrene tolerance studies, HEK293 cell suspensions were dispensed into 384-well microtiter plates at densities ranging from 500, 1000, to 2,000 cells per well in 40 µL growth medium. The assay plates contained 5 µL of polybrene in a doubling dilution series at a final concentration between 12 ng/mL and 25 µg/mL. At 96h post-seeding, cells were fixed with 4% PFA (w/v) for 20 minutes and stained for nuclei with solution containing 0.05% Triton X-100 (v/v) with 10 µM Hoechst for 10 minutes without any wash steps. Plates were imaged on the IN CELL Analyzer 2000 (INCA2000, GE Healthcare, USA) using a 4X magnifying objective allowing for one image per well, covering 100% of the well and analyzed using Developer 1.7 for Hoechst stained nuclei (NUCL).

#### Puromycin Assessment Studies for shRNA Lentiviral Particle Assay Development

For the puromycin kill curve, HEK293 cells were dispensed into 384-well microtiter plates at 1,000 cells per well in 45 µL of growth medium containing 8 µg/mL of polybrene. After 96h, transduction media was aspirated and 45 µL of growth media was dispensed containing a doubling dilution series of puromycin at a final concentration between 12 ng/mL and 25 µg/mL. After 168h incubation, assay plates were fixed and Hoechst stained as described above for analysis.

#### Transduction Studies for shRNA Lentiviral Particle Assay Development

For transduction studies with control lentiviral particles, HEK293 cells were dispensed into 384-well microtiter plates at 1,000 cells per well in 40 µL of growth media and settled overnight. Next, 10 µL of growth medium containing polybrene was dispensed into the assay plate at a final concentration of 8 µg/ml followed by transduction with control shRNA lentiviral particles. Three control lentiviral particles were used: 1) lentiviral particles containing scrambled shRNA (Sigma-Aldrich, Catalog #SHC002V); 2) lentiviral particles containing TurboGFP (Sigma-Aldrich, Catalog #SHC003V); and 3) lentiviral particles containing shRNA directed against PLK1 (Sigma-Aldrich, TRCN #0000121325). The control particles were diluted from stock into Dulbecco’s Modified Eagle’s Medium (DMEM) at 1×10^6^ transducing units (TU) per mL and 5 µL was transferred to the assay plates at a multiplicity of infection (MOI) of 5. The assay plates were centrifuged for 8 minutes at 1,300 rpm and incubated for 96h. Next, the transduction media was aspirated and 45 µL of growth media containing 1 µg/mL of puromycin was dispensed to the assay plate. After 168h incubation, assay plates were fixed and Hoechst stained as described above. Plates were imaged on the INCA2000 and analyzed using Developer 1.7 for NUCL and TurboGFP expression.

#### Image Acquisition and Analysis for Assay Development

Image acquisition for assay development was performed on the IN CELL Analyzer 2000 (INCA2000, GE Healthcare, USA) wide-field automated equipped with a large-chip CCD camera and custom-made polychroic with 4X magnifying objective allowing for whole well imaging. Images acquisition was captured at the following wavelengths: 350nm excitation / 455nm emission for Hoechst stained nuclei with an exposure time of 150 milli-seconds and 490nm excitation / 525nm emission for TurboGFP with an exposure time of 350 milli-seconds. One tile was imaged per well covering 100% of the well and required 5s per well with total imaging time of 35 minutes for a complete 384-well microtiter plate. Images were analyzed using the Developer 1.7 using the software’s built in segmentation algorithm. The segmentation algorithm determines NUCL and TurboGFP count based on the number of objects with pixel intensities above background and required a total analysis time of 15 minutes for a complete 384-well microtiter plate.

#### Genome-wide Arrayed shRNA Lentiviral Particle Library

The shRNA lentiviral particle library used for screening was custom made and purchased from Sigma-Aldrich. The shRNA lentiviral particle covers the whole human genome of 16,039 genes with approximately five shRNA clones per gene target including the 3’ UTR. The library contains 80,598 shRNA lentiviral particles arrayed as single clones into 295 384-well polypropylene source plates. Each 384-well source plate contains 18 μL of ready-to-use shRNA lentiviral particles at 1×10^6^ transducing units (TU) per mL. Columns 13-14 were left empty for high and low controls as set by the assay. As an additional quality control, rows O-P and columns 15-24 were empty for internal reference. The shRNA lentiviral particle library was stored at - 80°C until use.

### Supplementary Figure and Table legends

**Supplementary Figure 1.**
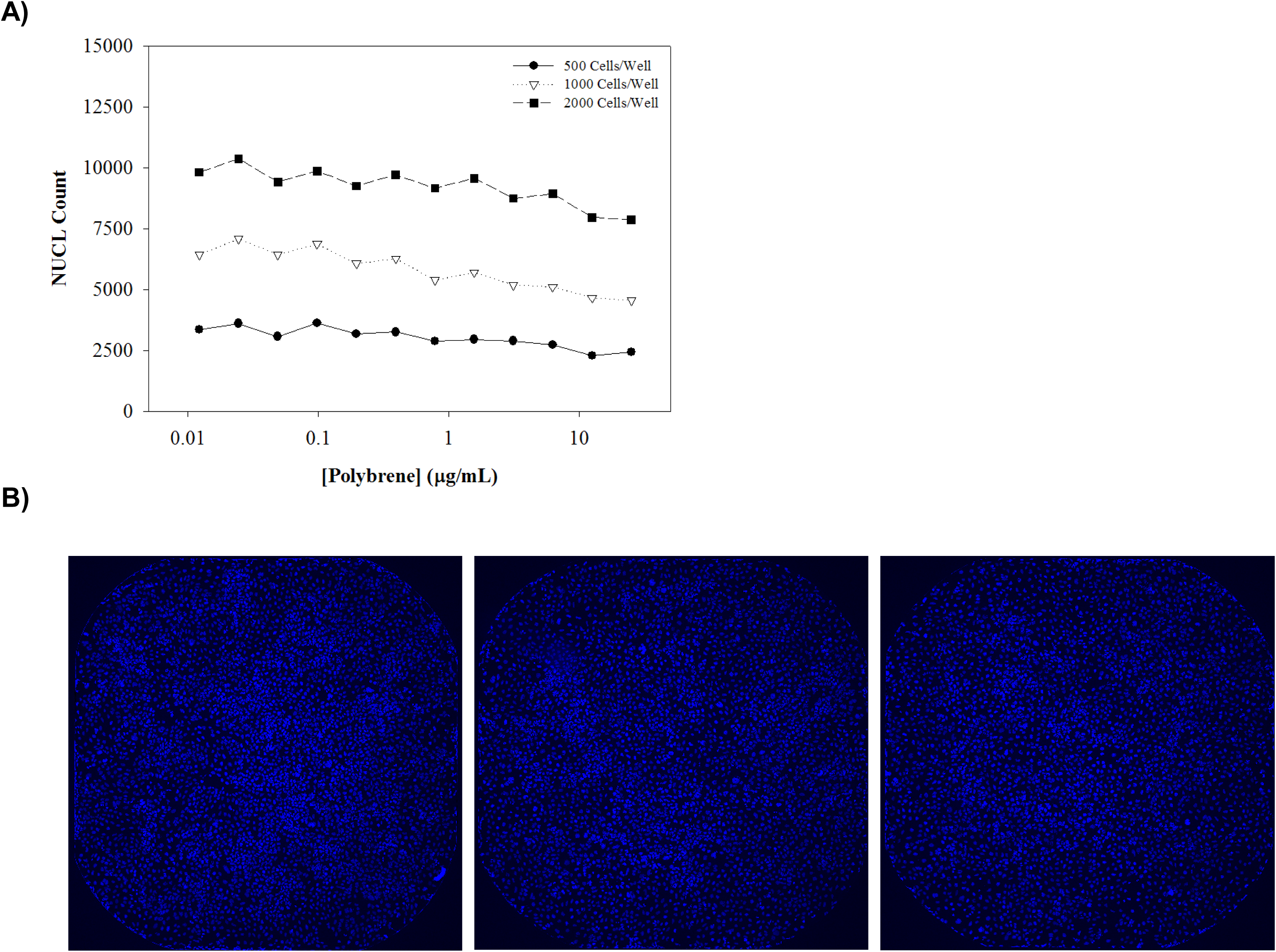
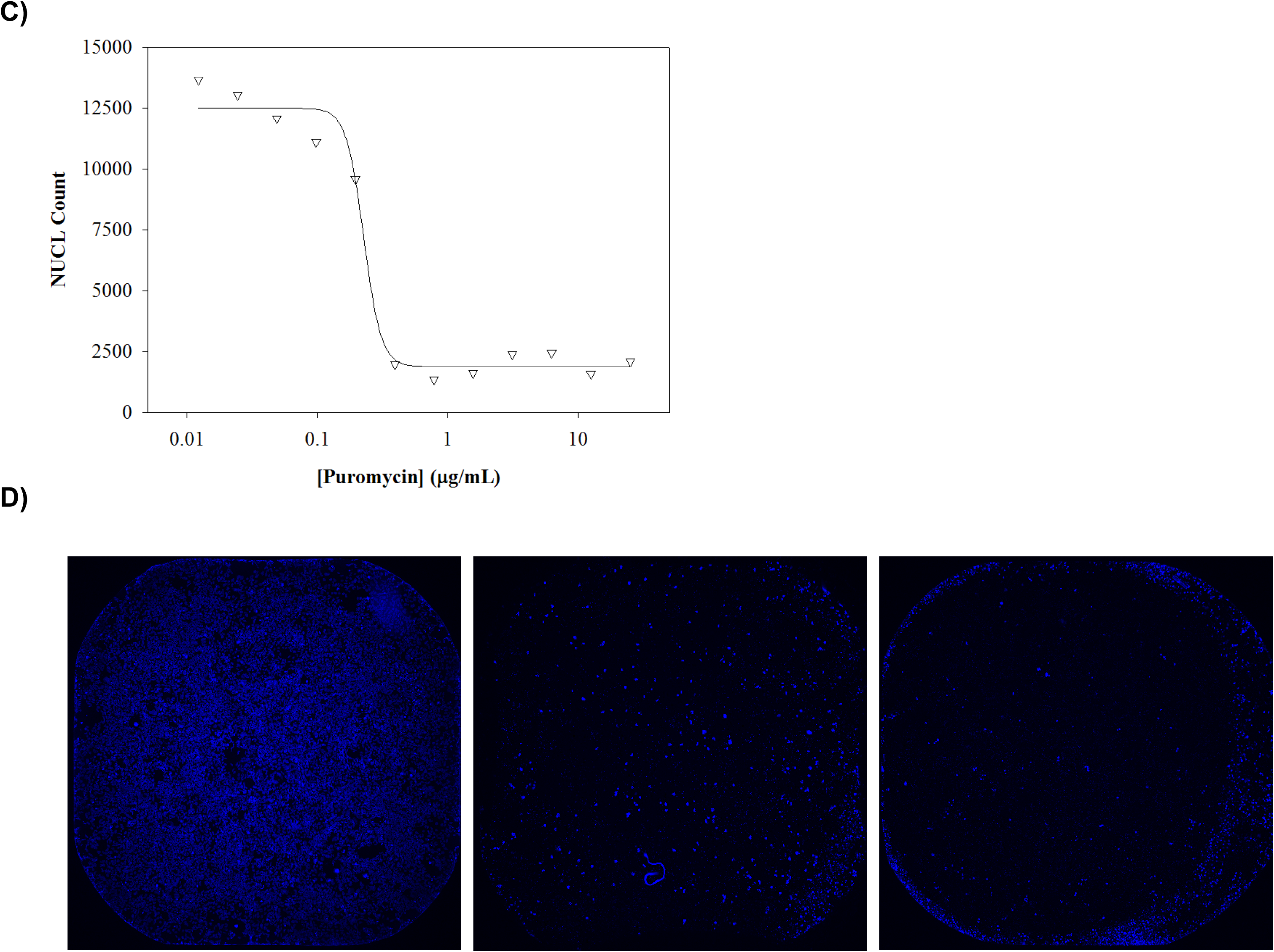
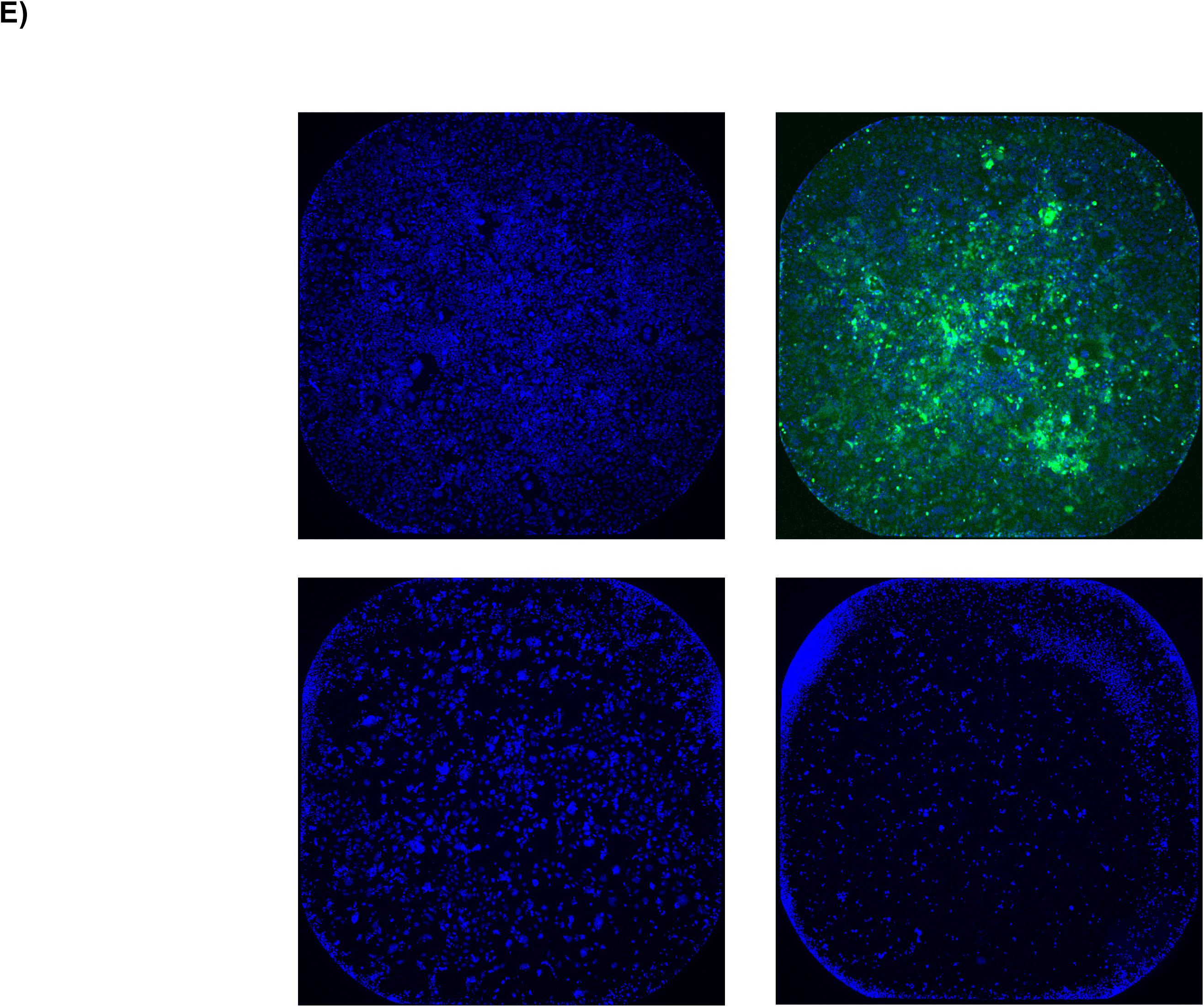
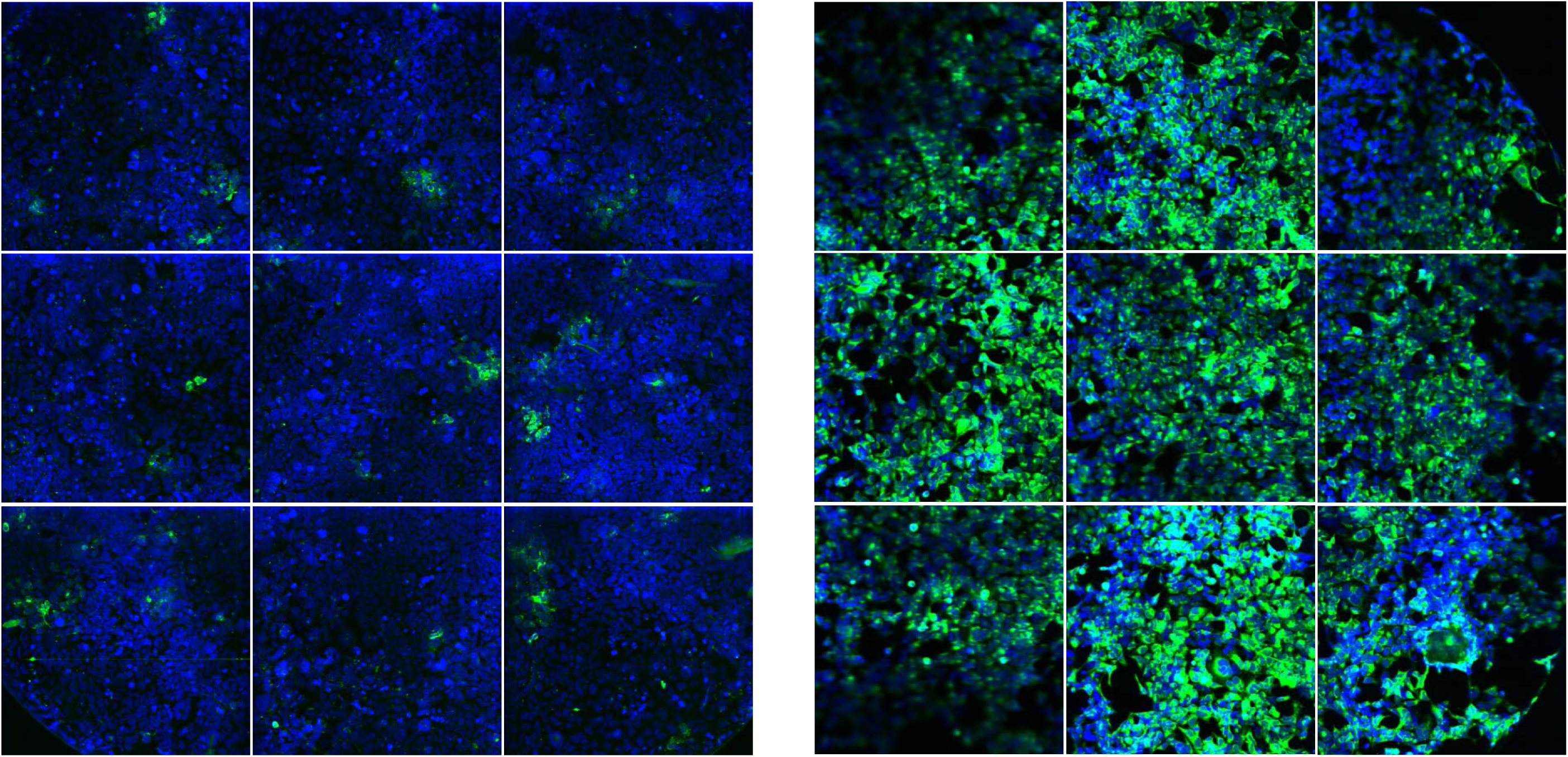
Assay development for shRNA lentiviral transduction. Development studies in 384-well microtiter plate format of DENV shRNA infection assay. **A**) HEK293 cell were seeded in cell densities ranging from 500 (●), 1000 (▼), and 2000 (■) cells per well in the presence of polybrene at 12-point doubling concentration series between 12 ng/mL and 25 μg/mL. For each measurement, averages are calculated from 3 data points. **B**) Images of Hoechst stained nuclei at 96h post seeding for 1,000 cells per well in polybrene: Left image at 0.01 μg/mL; Middle image at 6.25 μg/mL; and Right image at 25 μg/mL. Images were acquired using the INCA2000 with blue channel for detection of Hoechst stained nuclei. **C**) HEK293 cell were seeded at 1,000 (▼) cells per well in the presence of polybrene at 8 μg/mL for 96h. Next, cells were incubated with puromycin in a 12-point doubling concentration series between 12 ng/mL and 25 μg/mL for 168h. For each measurement, averages are calculated from 3 data points. D) Images of Hoechst stained nuclei at 168h post puromycin incubation: Left image at 0.01 μg/mL; Middle image at 0.78 μg/mL; and Right image at 25 μg/mL. Images were acquired using the INCA2000 with blue channel for detection of Hoechst stained nuclei. E) HEK293 cell were seeded at 1,000 cells per well in the presence of polybrene at 8 μg/mL followed by transduction with control lentiviral particles for 96h. Next, positive transduced cells were selected at 1 μg/mL and images at 168h post puromycin incubation: Top left image of lentiviral particle containing scrambled shRNA; Top right image of lentiviral particle containing TurboGFP; Bottom left image of lentiviral particle containing shRNA targeting PLK1; and Bottom right image of untransduced well. Images were acquired using the INCA2000 with blue channel for detection of Hoechst stained nuclei and green channel for detection of TurboGFP.

**Supplementary Figure 2.**
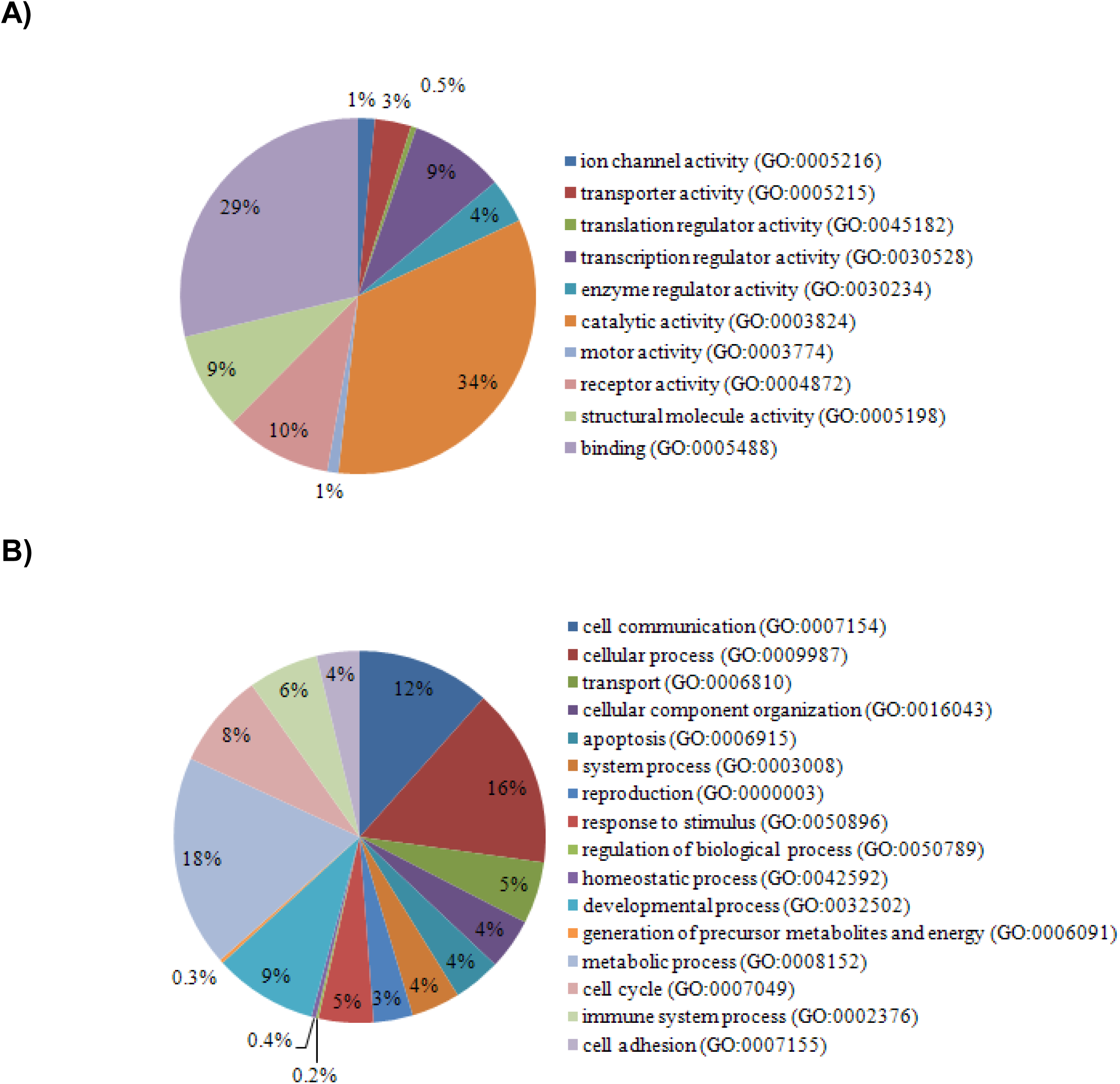
Classification of the 313 common inhibitors into **A)** molecular and **B)** biological functional categories as obtained from Panther classification system. The relative percentage of the genes involved in each category is represented in the pie charts.

**Supplementary Figure 3.**
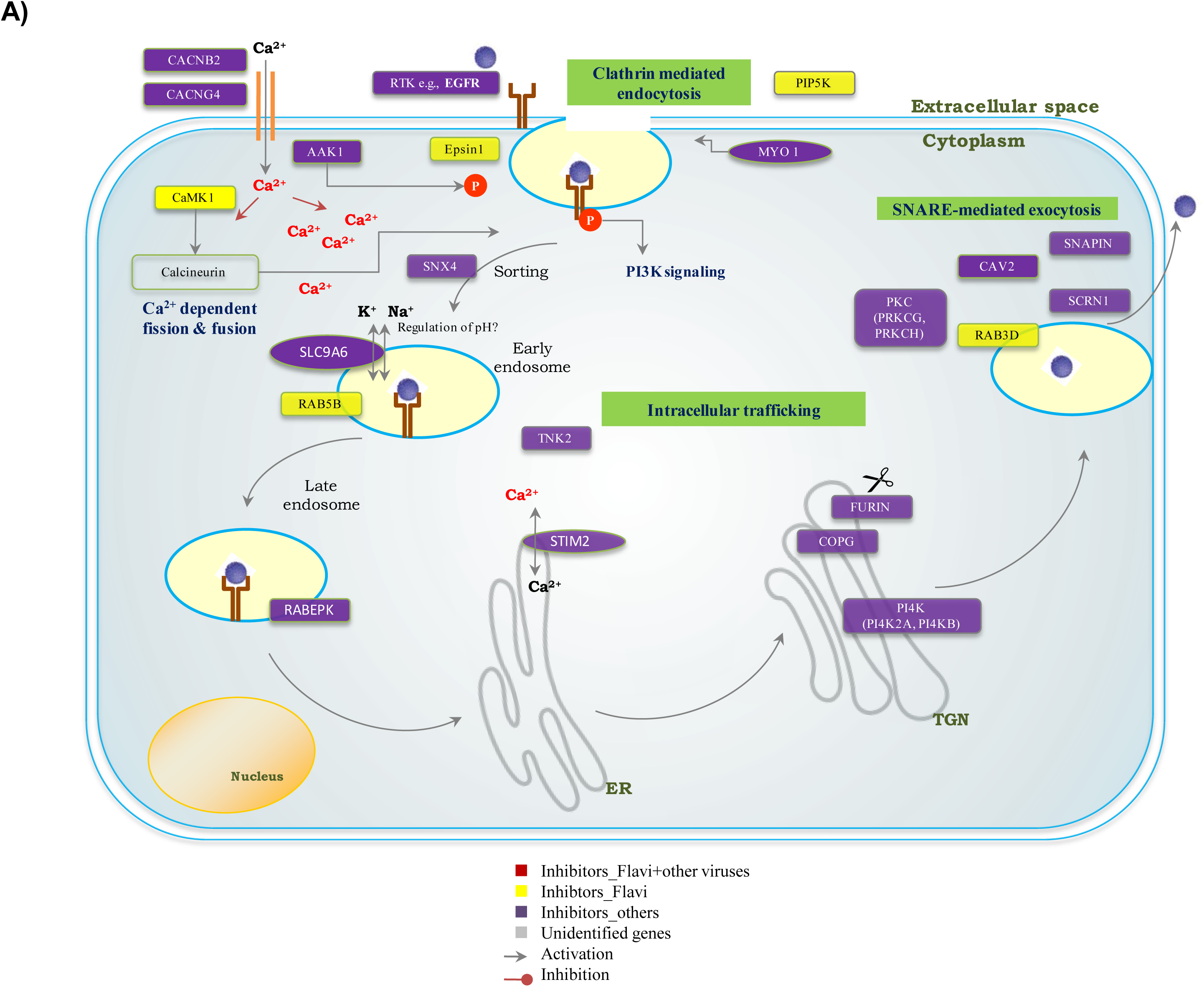
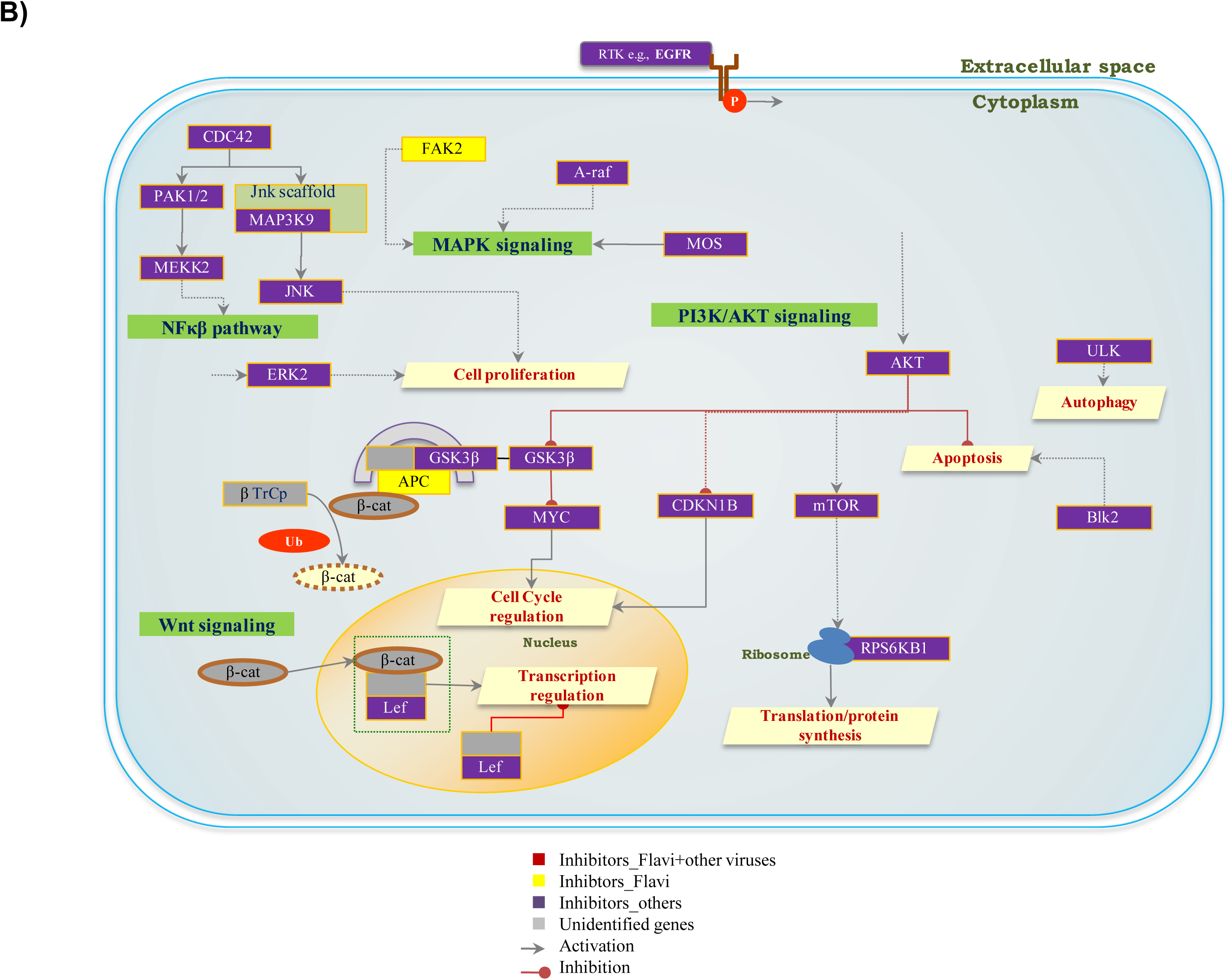
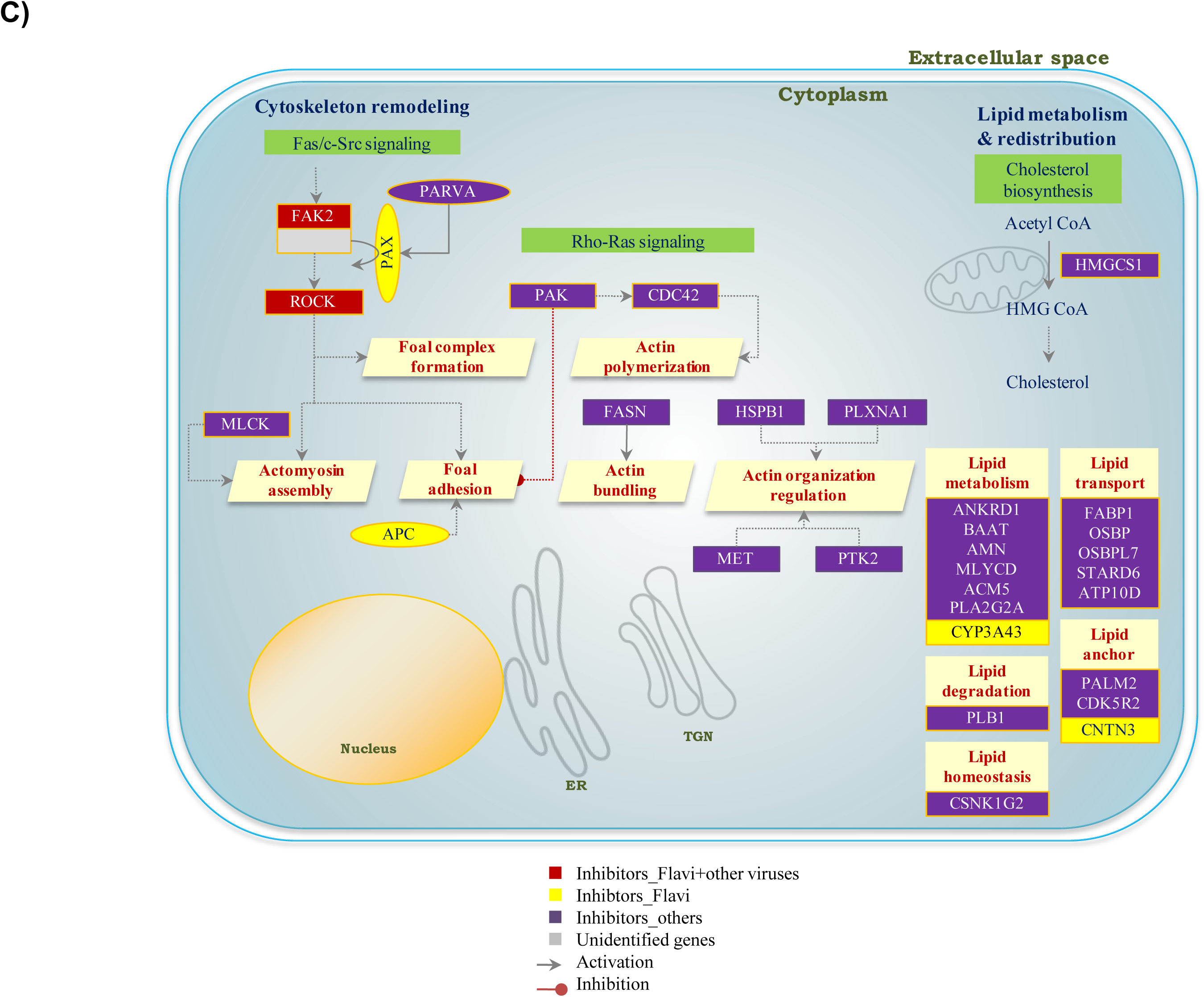
Pathways and processes plausibly modulated or perturbed due to host cellular machinery hijack by the virus. **A)** Clathrin mediated endocytosis, intracellular trafficking and exocytosis. 2 interactions not found in canonical clathrin-mediated endocytosis are also shown – AAK1 regulation of AP2 complex (Conner SD, Schmid SL: Identification of an adaptor-associated kinase, AAK1, as a regulator of clathrin-mediated endocytosis. J Cell Biol. 2002 ;156(5):921-9.), MYO1 in vesicle pinching (Krendel M, Osterweil EK, Mooseker MS: Myosin 1E interacts with synaptojanin-1 and dynamin and is involved in endocytosis. FEBS Lett. 2007 ;581(4):644-50.) **B)** Signalling pathways including MAPK signaling cascade, PI3/AKT signaling, Wnt-signaling and NFkb pathway. **C)** Cyotoskeleton remodeling and lipid metabolism.

**Supplementary Table 1.**
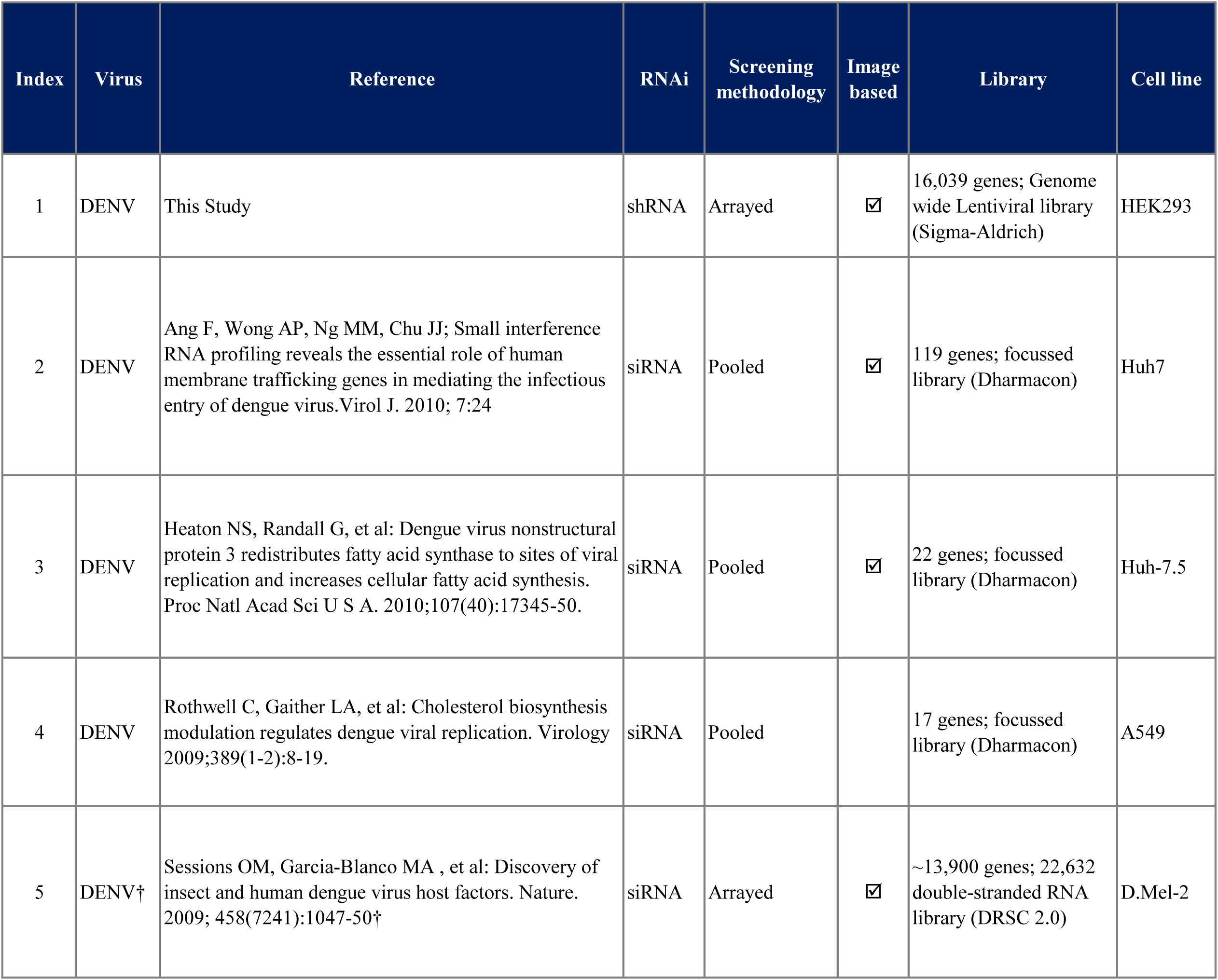
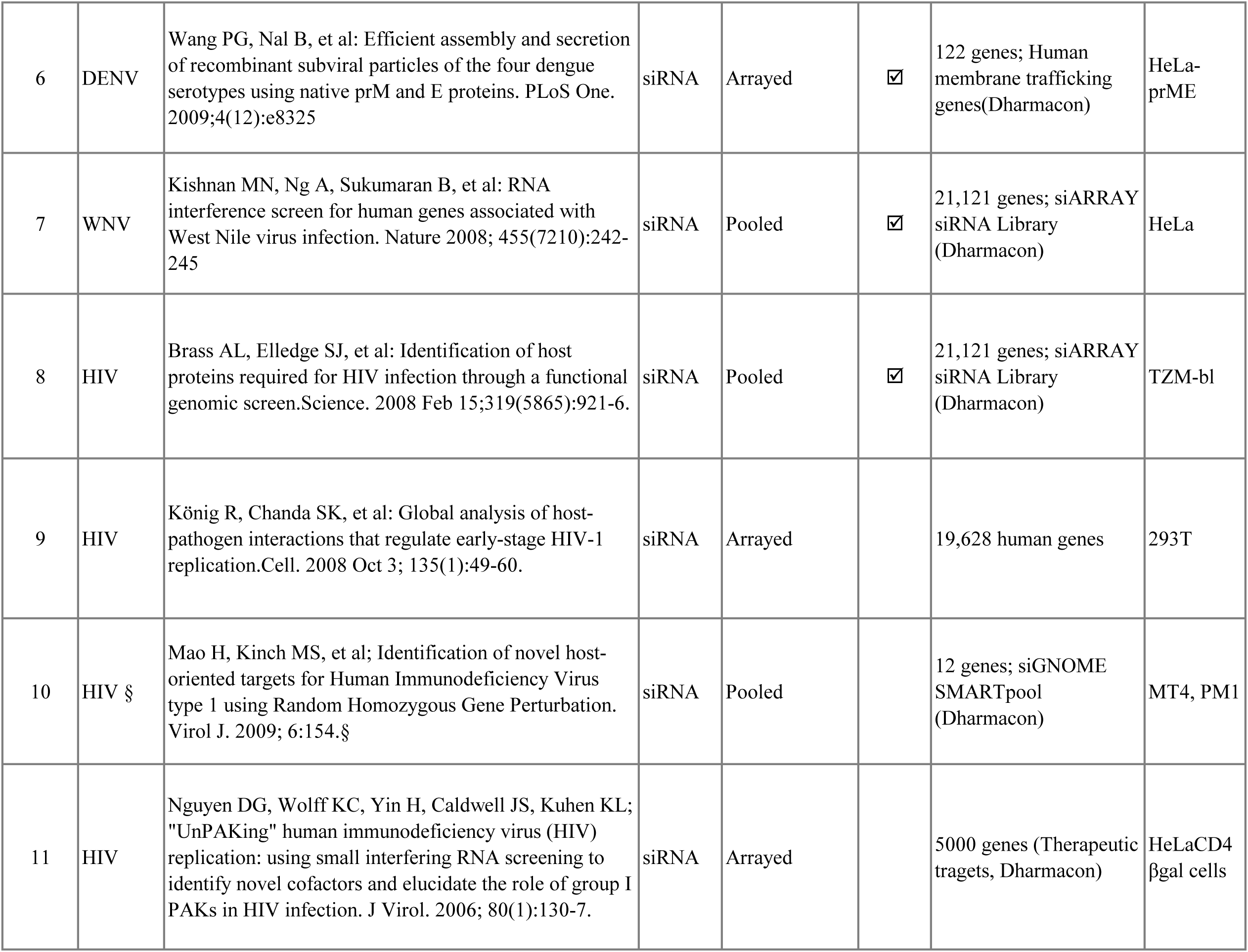
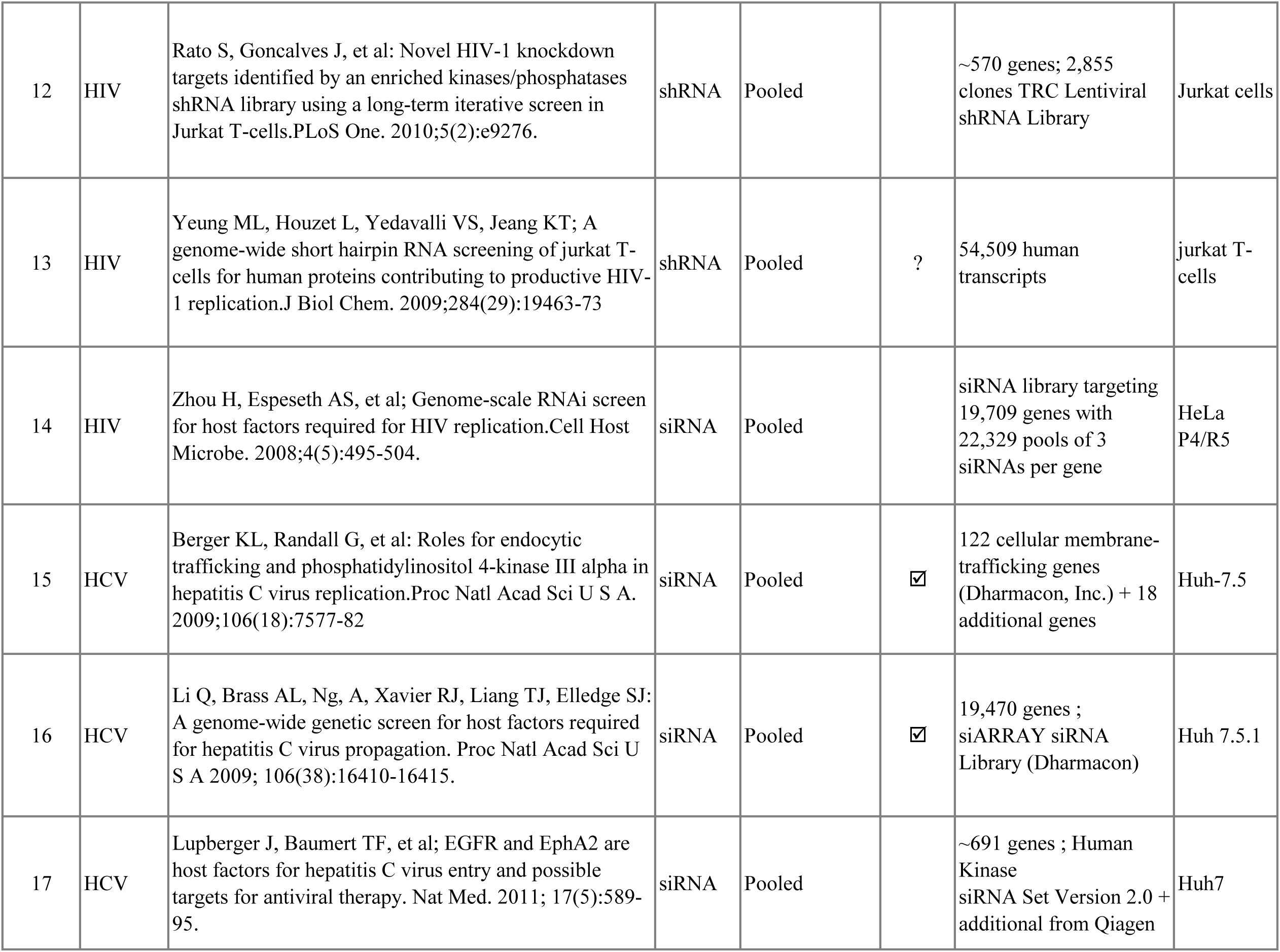
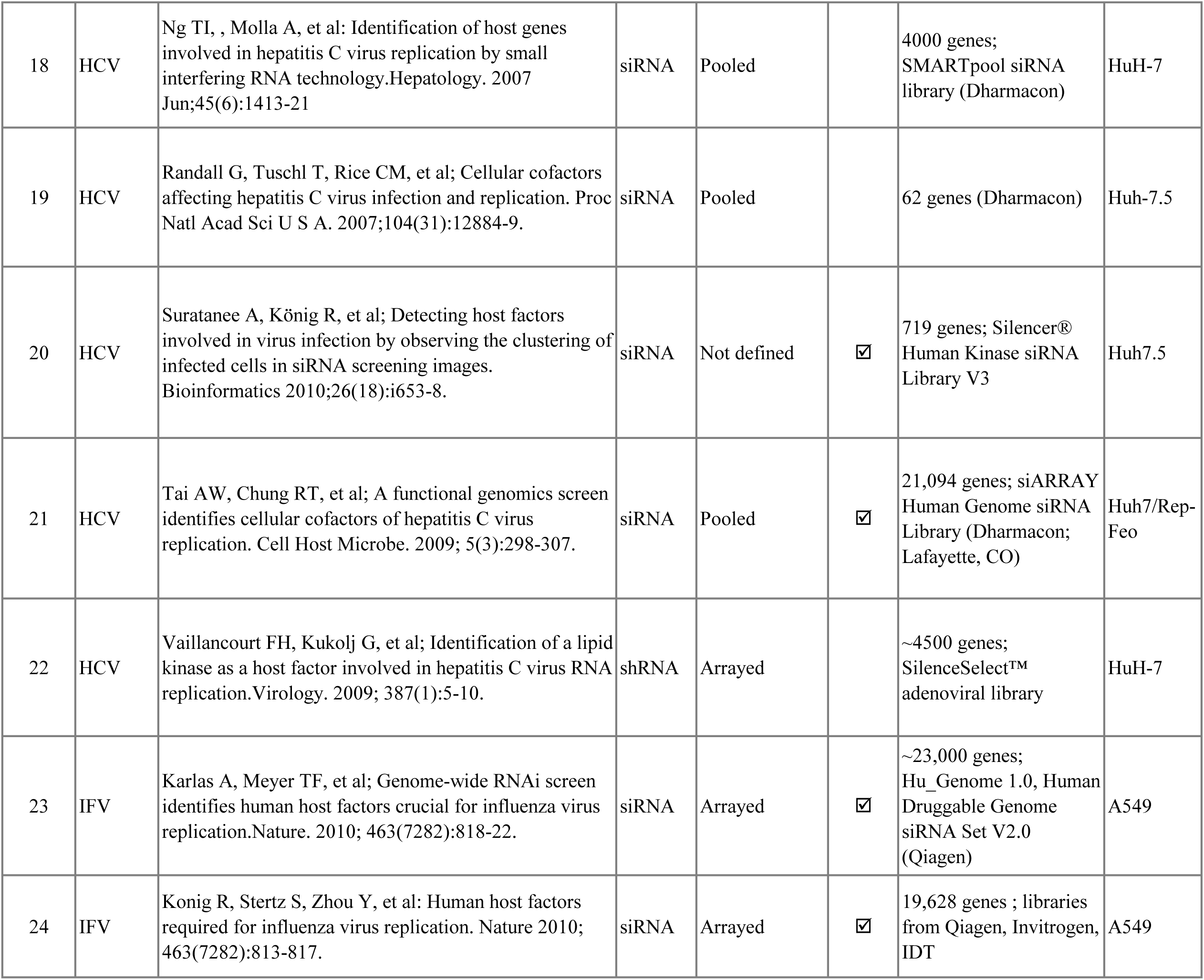
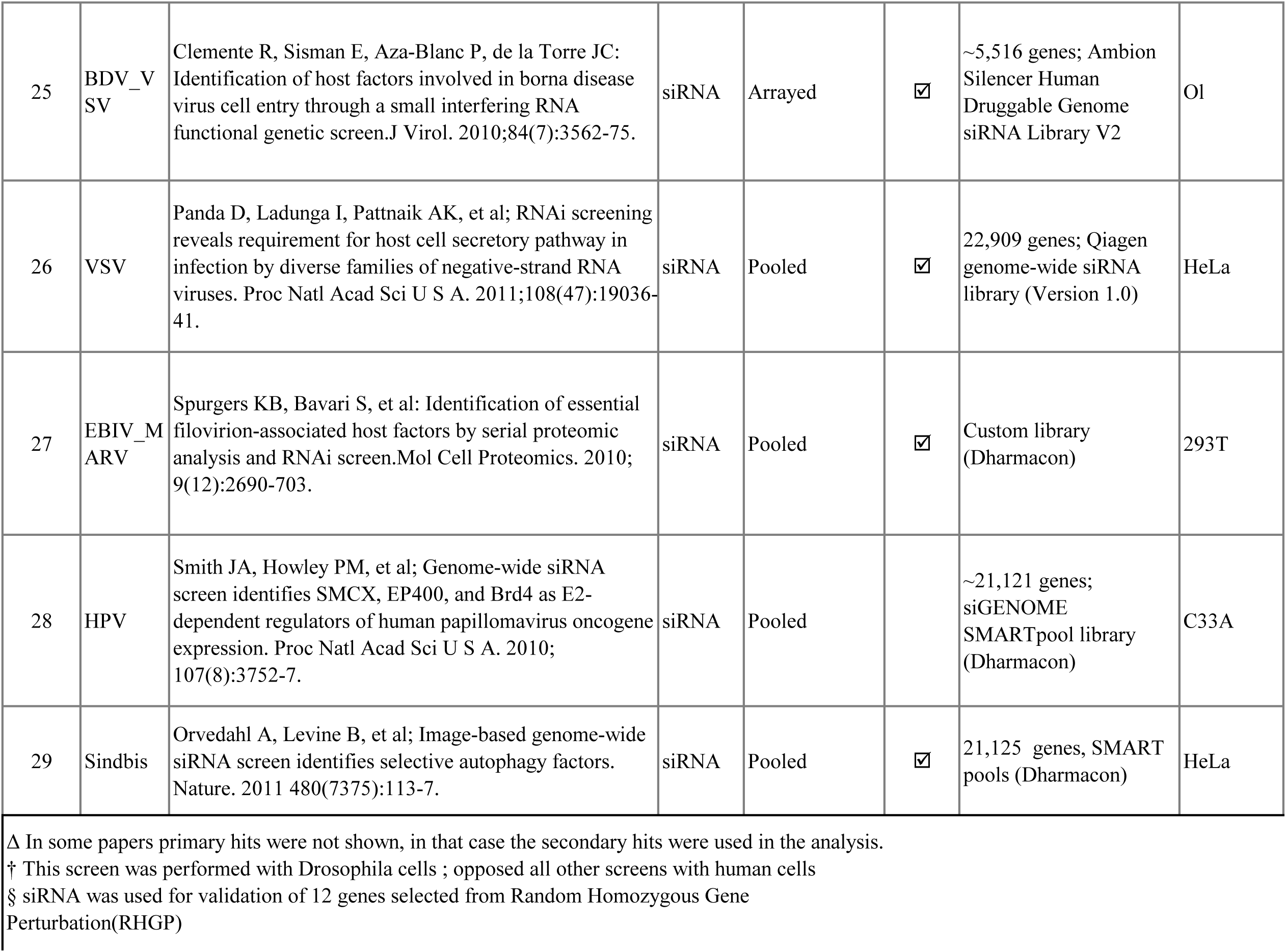
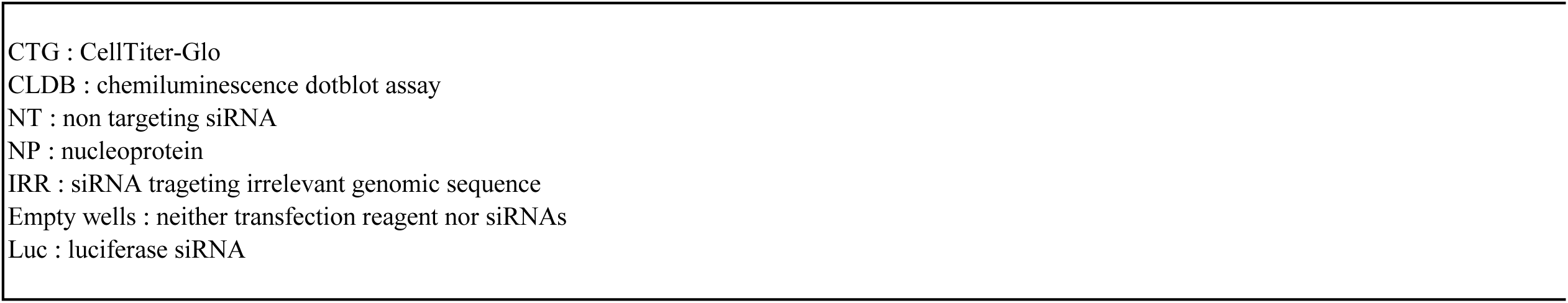
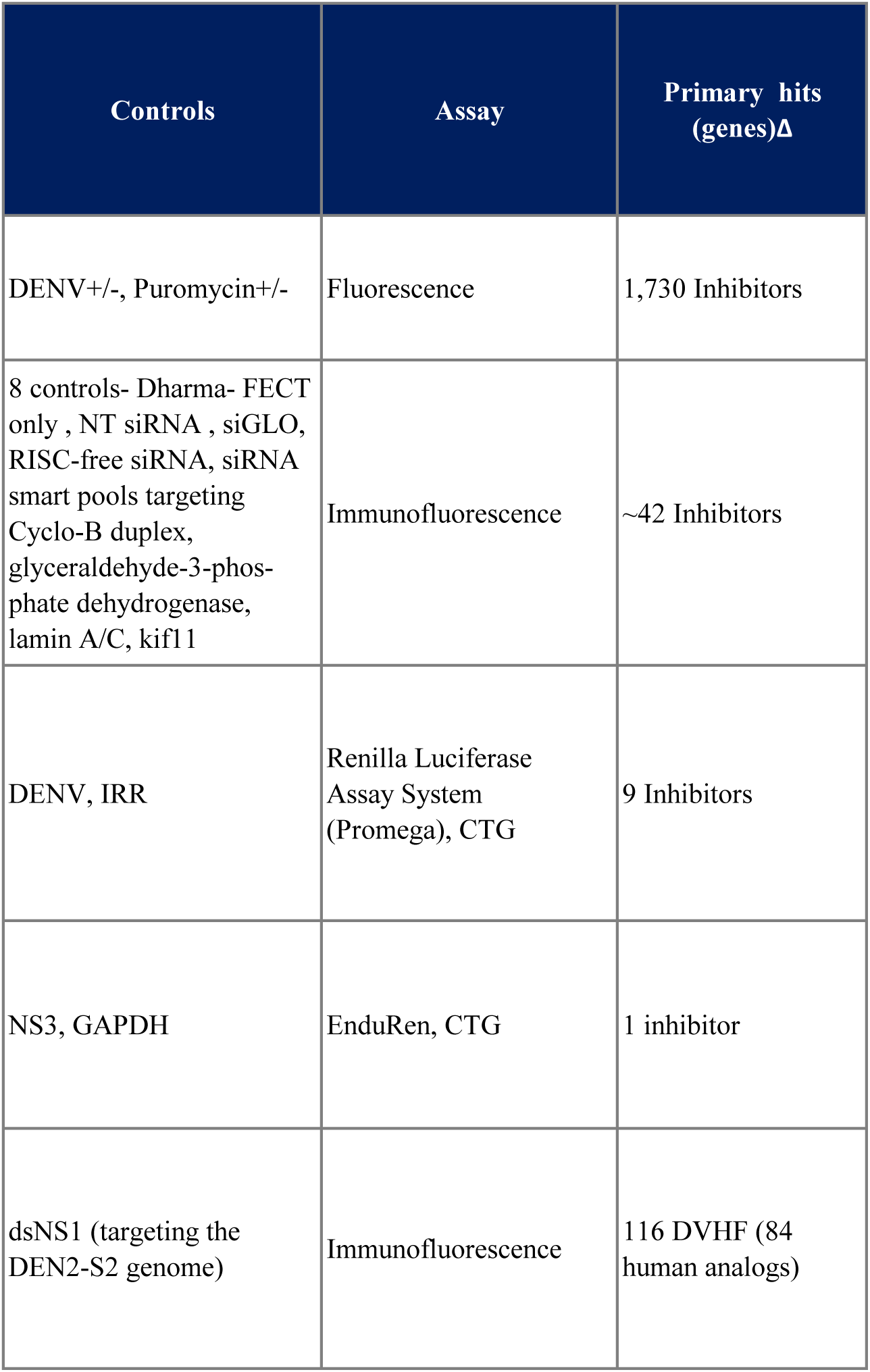
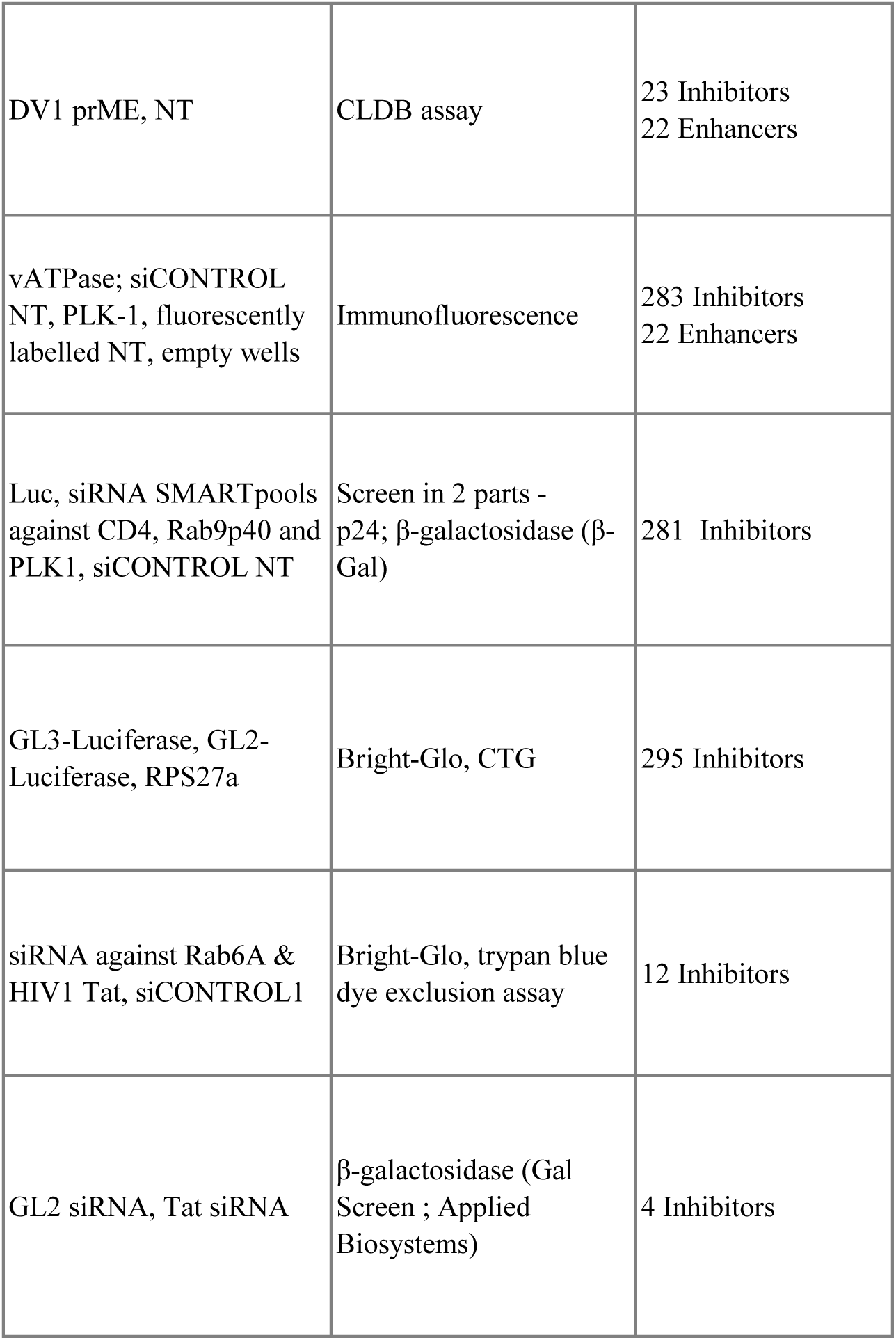
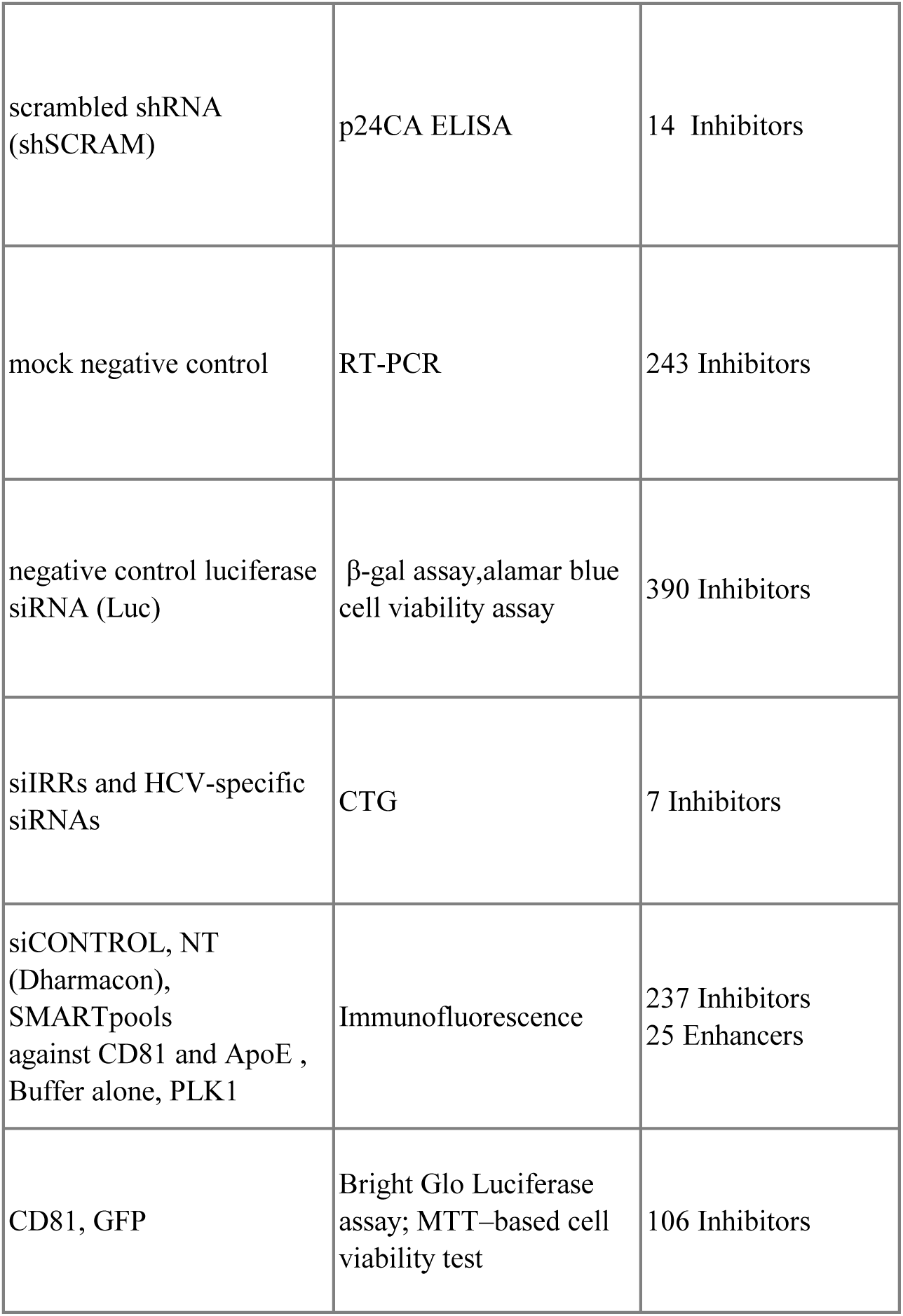
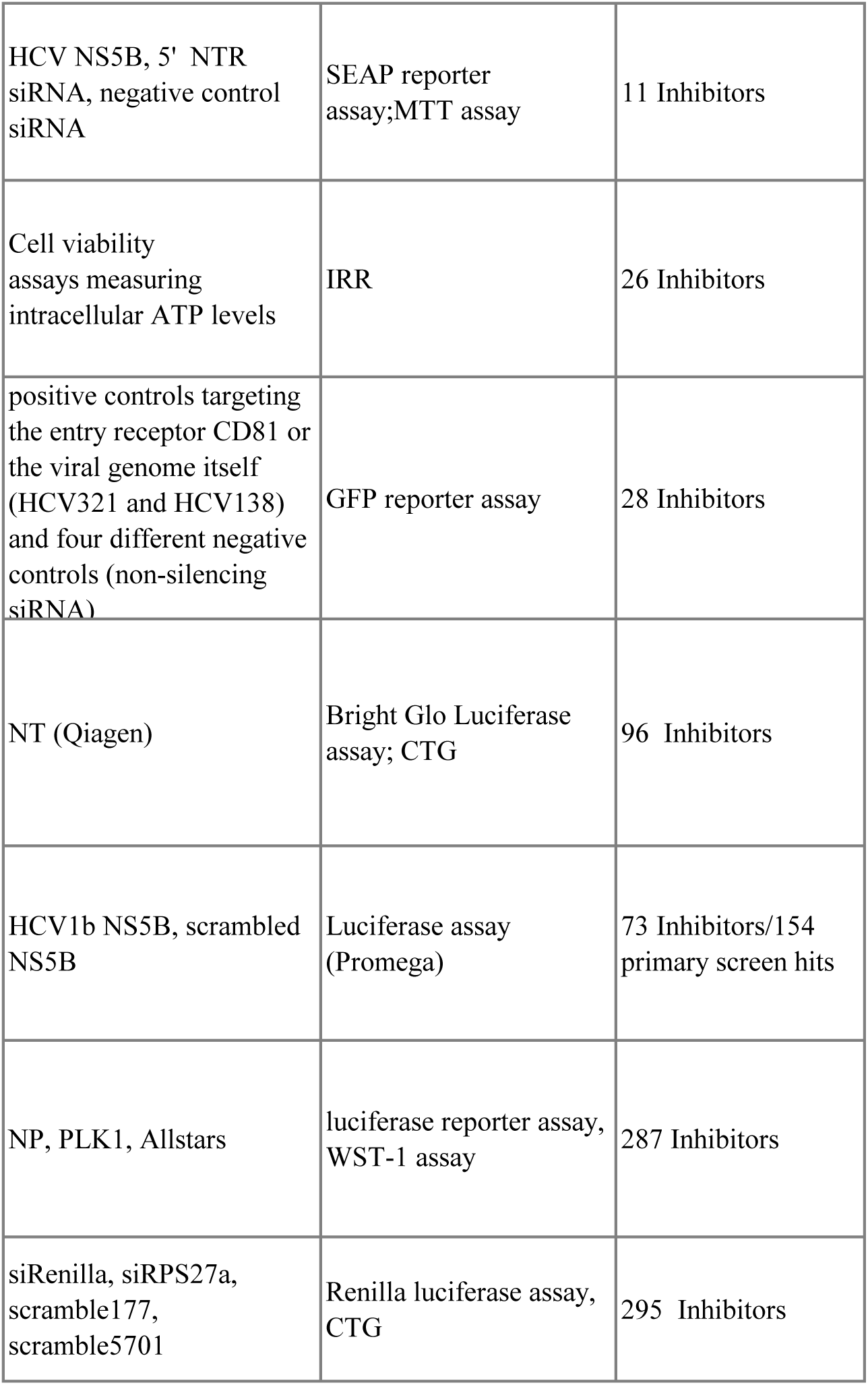
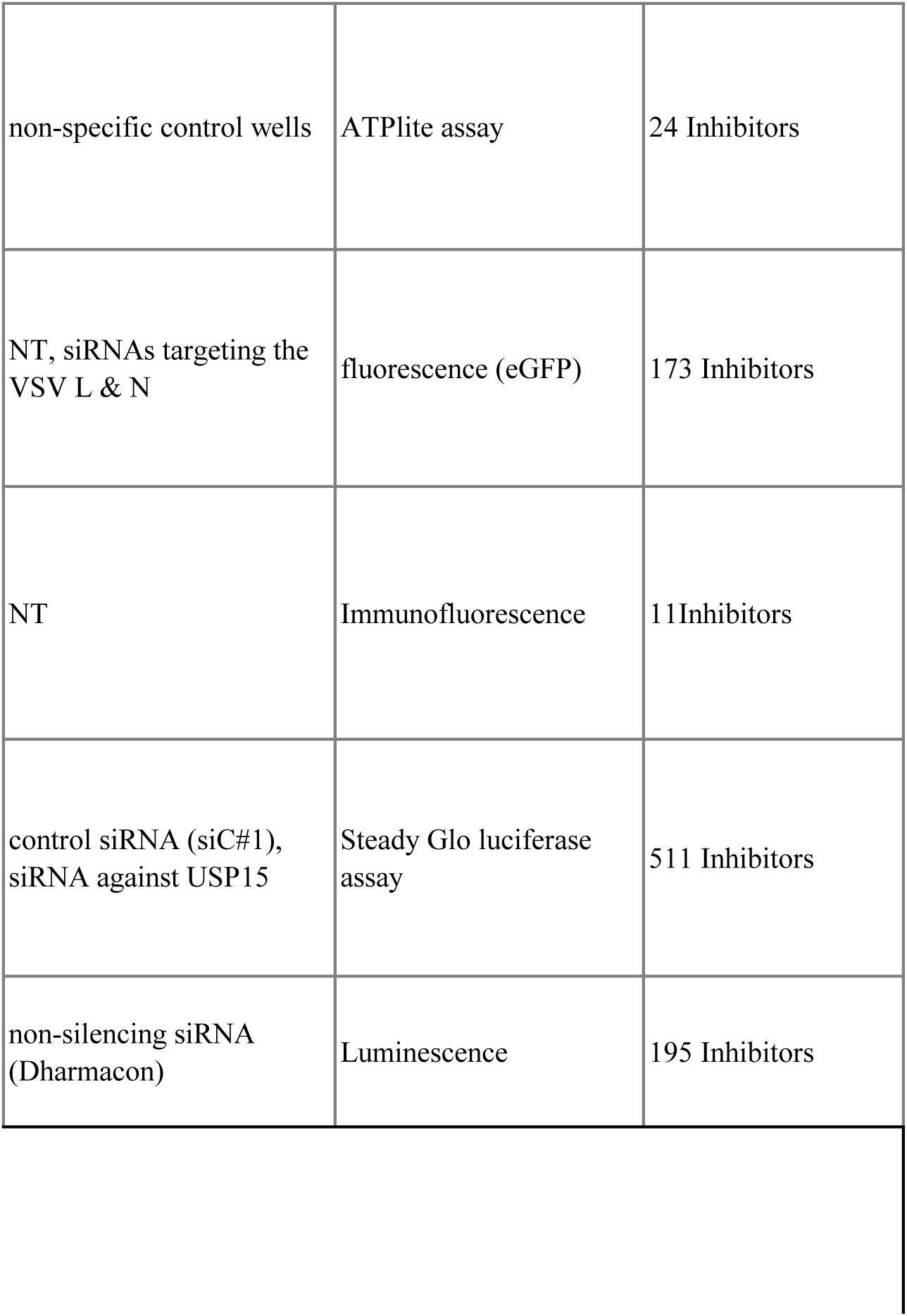
A summary of the 28 RNAi screening papers used to obtain the published set of host-viral factors across different viruses. The methodologies and the number of hits reported in each study are shown in this table.

**Supplementary Table 2.** List of 40 inhibitors that were common between the flavivirus RNAi screening publications and the screen reported in this paper.

| Gene | Virus | NCBI_ID | Gene Description |
| --- | --- | --- | --- |
| RSPO3 | Flavivirus | NM_032784 | R-spondin 3 homolog (*Xenopus laevis*) |
| DEFB129 | Flavivirus | NM_080831 | defensin, beta 129 |
| MUC2 | Flavivirus | NM_002457 | Homo sapiens mucin 2, oligomeric mucus/gel-forming (MUC2), mRNA |
| SIRPA | Flavivirus | NM_080792 | Homo sapiens signal-regulatory protein alpha (SIRPA), transcript variant 3, mRNA |
| PCDHGA10 | Flavivirus | NM_018913 | protocadherin gamma subfamily A, 10 |
| CNTN3 | Flavivirus | NM_020872 | contactin 3 (plasmacytoma associated) |
| CSNK2A1 | Flavivirus + Other_viruses | NM_001895/<br>NM_177559 | Homo sapiens casein kinase 2, alpha 1 polypeptide (CSNK2A1), transcript variant 2, mRNA/, transcript variant 1, mRNA |
| TNFSF13B | Flavivirus + Other_viruses | NM_006573 | Homo sapiens tumor necrosis factor (ligand) superfamily, member 13b (TNFSF13B), mRNA |
| EPN1 | Flavivirus | NM_013333 | Homo sapiens epsin 1 (EPN1), transcript variant 3, mRNA |
| RAB5B | Flavivirus | NM_002868 | Homo sapiens RAB5B, member RAS oncogene family (RAB5B), mRNA |
| CFHR5 | Flavivirus + Other_viruses | NM_030787 | complement factor H-related 5 |
| CAMK1 | Flavivirus | NM_003656 | Homo sapiens calcium/calmodulin-dependent protein kinase I (CAMK1), mRNA |
| PTK2B | Flavivirus + Other_viruses | NM_004103 | Homo sapiens PTK2B protein tyrosine kinase 2 beta (PTK2B), transcript variant 2, mRNA |
| KATNAL1 | Flavivirus | NM_032116 | katanin p60 subunit A-like 1 |
| PXN | Flavivirus | NM_002859 | paxillin |
| APC | Flavivirus | NM_000038 | Homo sapiens adenomatous polyposis coli (APC), transcript variant 3, mRNA |
| ROCK1 | Flavivirus + Other_viruses | NM_005406 | Homo sapiens Rho-associated, coiled-coil containing protein kinase 1 (ROCK1), mRNA |
| ROCK2 | Flavivirus + Other_viruses | NM_004850 | Homo sapiens Rho-associated, coiled-coil containing protein kinase 2 (ROCK2), mRNA |
| WDR18 | Flavivirus + Other_viruses | NM_024100 | Homo sapiens WD repeat domain 18 (WDR18), mRNA |
| TRIM63 | Flavivirus | NM_032588 | Homo sapiens tripartite motif-containing 63 (TRIM63), mRNA |
| NDUFB3 | Flavivirus | NM_002491 | NADH dehydrogenase (ubiquinone) 1 beta subcomplex, 3, 12kDa |
| RAB3D | Flavivirus | NM_004283 | Homo sapiens RAB3D, member RAS oncogene family (RAB3D), mRNA |
| PLK4 | Flavivirus + Other_viruses | NM_014264 | Homo sapiens polo-like kinase 4 (Drosophila) (PLK4), mRNA |
| PAXIP1 | Flavivirus | NM_007349 | PAX interacting (with transcription-activation domain) protein 1 |
| CHML | Flavivirus | NM_001821 | choroideremia-like (Rab escort protein 2) |
| GORASP1 | Flavivirus | NM_031899 | golgi reassembly stacking protein 1, 65kDa |
| NIPA2 | Flavivirus | NM_030922 | non imprinted in Prader-Willi/Angelman syndrome 2 |
| OAZ3 | Flavivirus | NM_016178 | ornithine decarboxylase antizyme 3 |
| PIP5K1A | Flavivirus | NM_003557 | Homo sapiens phosphatidylinositol-4-phosphate 5-kinase, type I, alpha (PIP5K1A), mRNA |
| NPR2 | Flavivirus | NM_000907 | Homo sapiens natriuretic peptide receptor B/guanylate cyclase B (atrionatriuretic peptide receptor B) (NPR2), transcript variant 1, |
| TAF4B | Flavivirus | XM_290809 | TAF4b RNA polymerase II, TATA box binding protein (TBP)-associated factor, 105kDa |
| TEAD1 | Flavivirus | NM_021961 | TEA domain family member 1 (SV40 transcriptional enhancer factor) |
| ZNF91 | Flavivirus | NM_003430 | zinc finger protein 91 |
| KIF17 | Flavivirus + Other_viruses | NM_020816 | Homo sapiens kinesin family member 17 (KIF17), transcript variant 1, mRNA |
| KLHL3 | Flavivirus | NM_017415 | kelch-like 3 (Drosophila) |
| GLCCI1 | Flavivirus + Other_viruses | NM_138426;<br>XM_379784 | glucocorticoid induced transcript 1 |
| KIAA1539 | Flavivirus | NM_025182 | KIAA1539 |
| TTC33 | Flavivirus | NM_012382 | tetratricopeptide repeat domain 33 |
| CALCB | Flavivirus + Other_viruses | NM_000728 | calcitonin-related polypeptide beta |
| CYP3A43 | Flavivirus | NM_022820 | Homo sapiens cytochrome P450, family 3, subfamily A, polypeptide 43 (CYP3A43), transcript variant 1, mRNA |

| Function |
| --- |
| Activator of the beta-catenin signaling cascade |
| Antimicrobial/ Beta-defensin |
| Antimicrobial/ Protective barrier against particles and infectious agents at mucosal surface |
| Blood coagulation |
| Cell adhesion |
| Cell adhesion |
| Cell cycle |
| Cell proliferation, immune response |
| Clathrin mediated endocytosis |
| Clathrin mediated endocytosis |
| Complement activation |
| component of a calmodulin-dependent protein kinase cascade |
| Cytoskeleton |
| Cytoskeleton |
| Cytoskeleton |
| Cytoskeleton |
| Cytoskeleton, apoptosis |
| Cytoskeleton, apoptosis |
| Development |
| E3 ubiquitin ligase |
| Electron transport |
| Exocytosis |
| G2/M transition |
| genome stability |
| Geranylgeranylation of Rab proteins |
| Golgi apparatus integrity |
| Ion transport |
| ornithine decarboxylase inhibitor activity |
| phosphatidylinositol phosphate kinase activity |
| Suppresses Src-dependent tyrosine phosphorylation and activation of PDPK1 |
| Transcription factor |
| Transcription factor |
| Transcription regulation |
| Transport, motor protein |
| ubiquitination and proteasomal degradation |
| uncharacterized |
| uncharacterized |
| uncharacterized |
| Vasodilation |
| Xenobiotic metabolic process |

**Supplementary Table 3.**
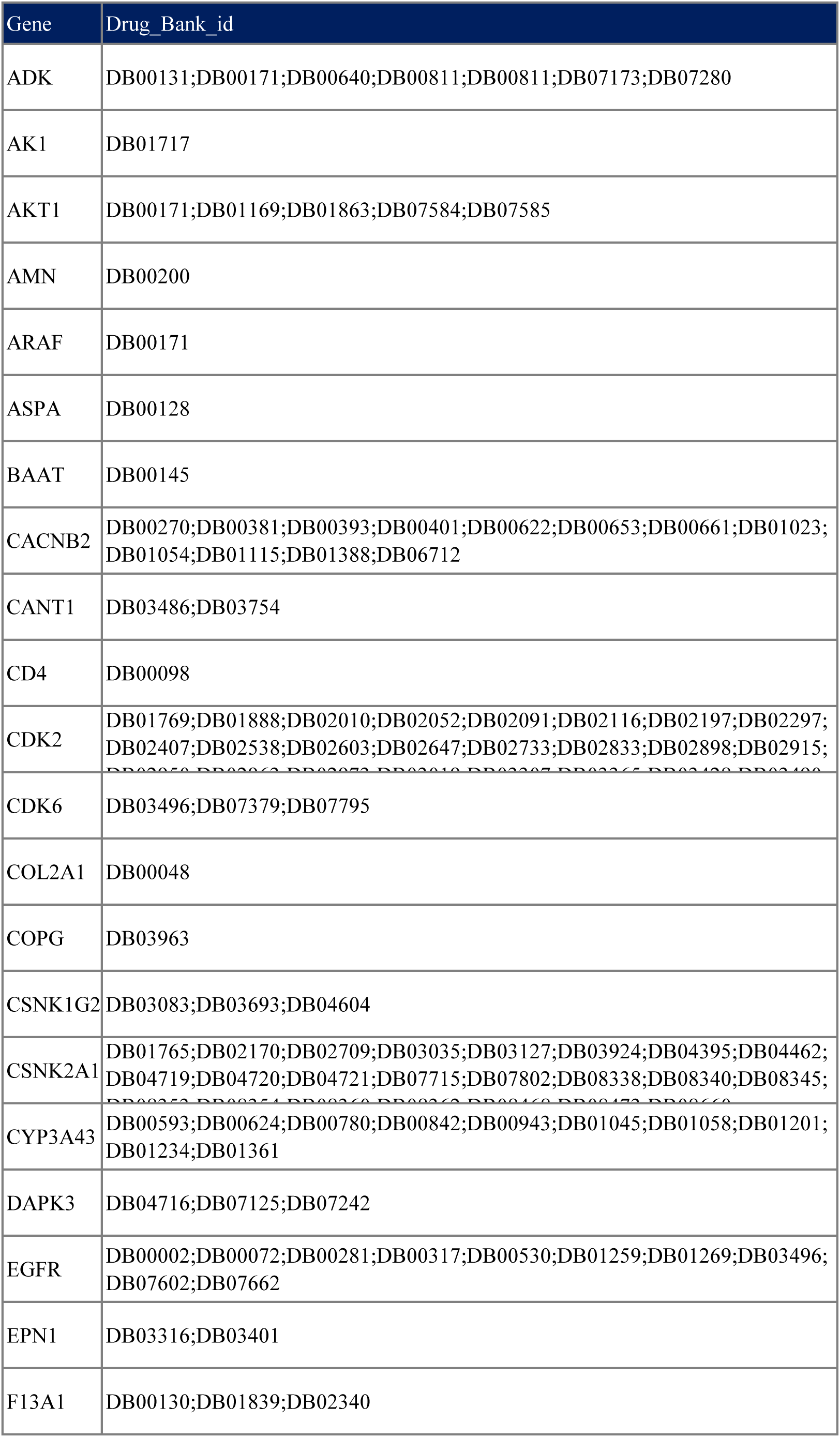
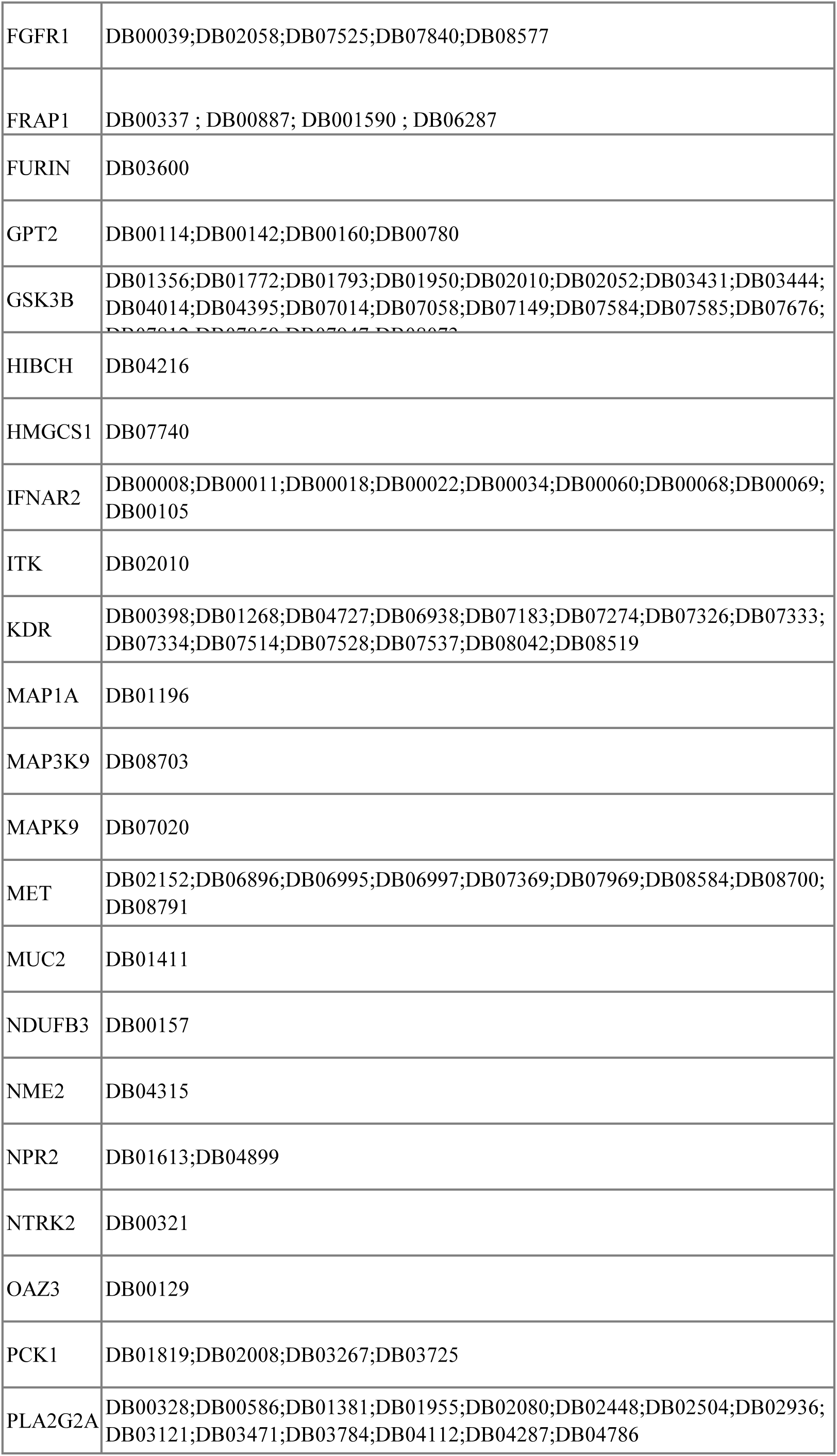
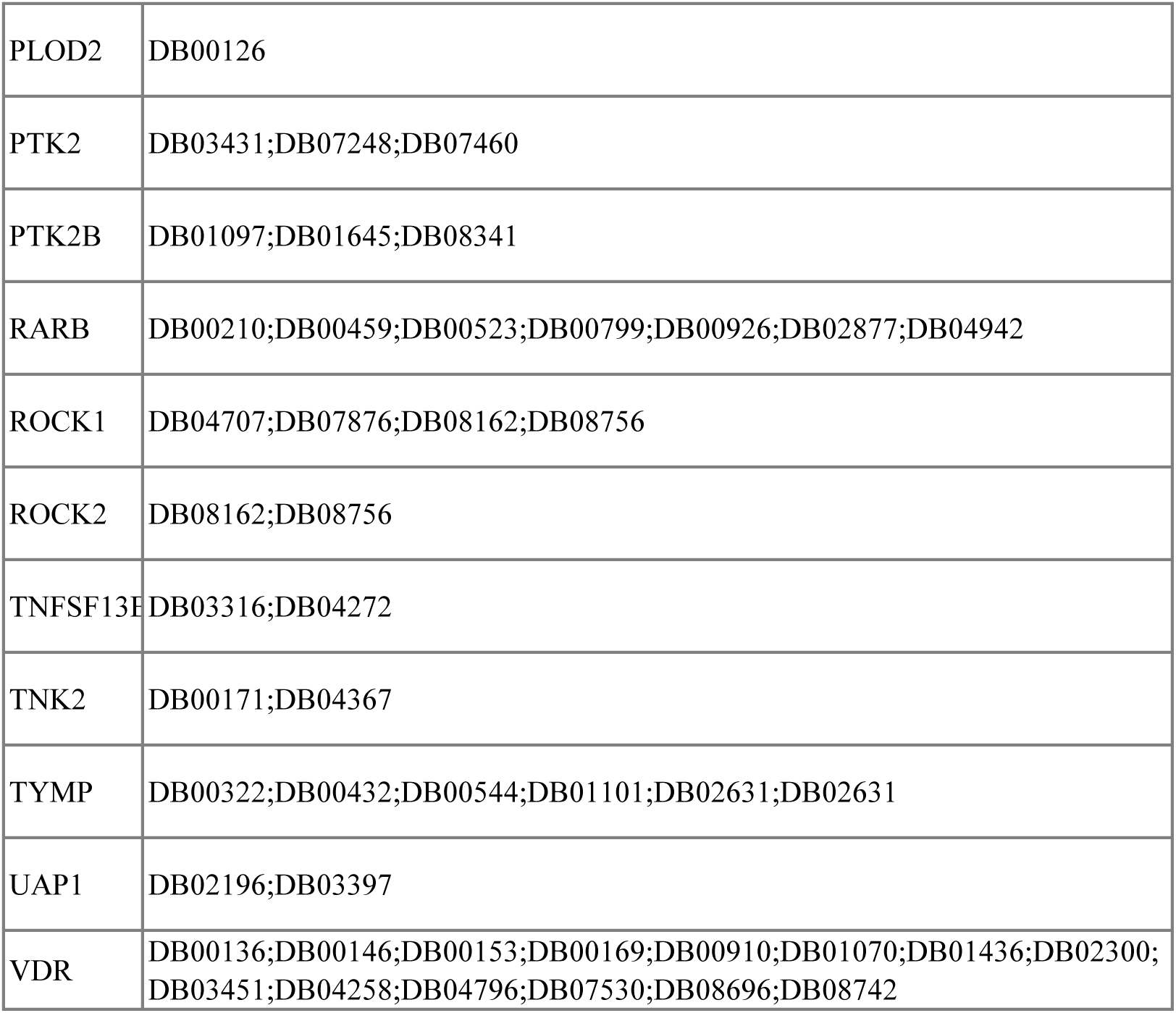
The Druggable targets along with their Drug bank identifiers are represented in this table. These targets are set of inhibitors obtained from 313 overlapping host viral factors.

**Supplementary Table 4.**
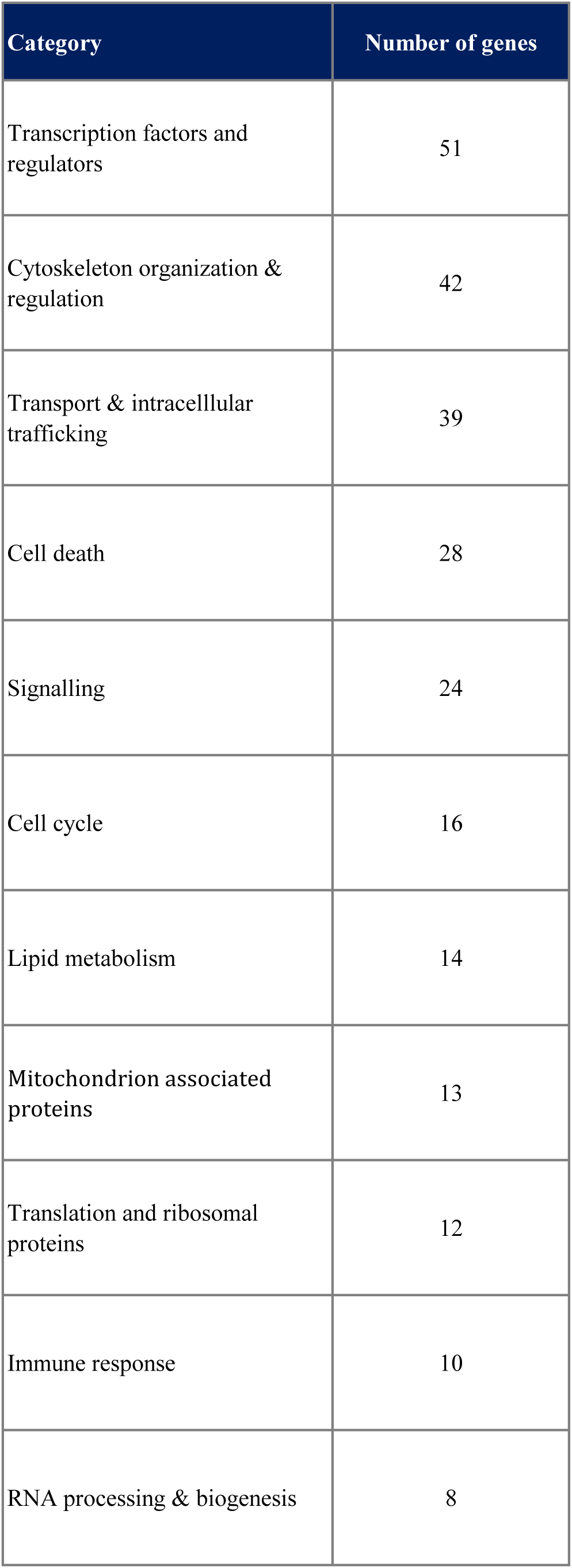
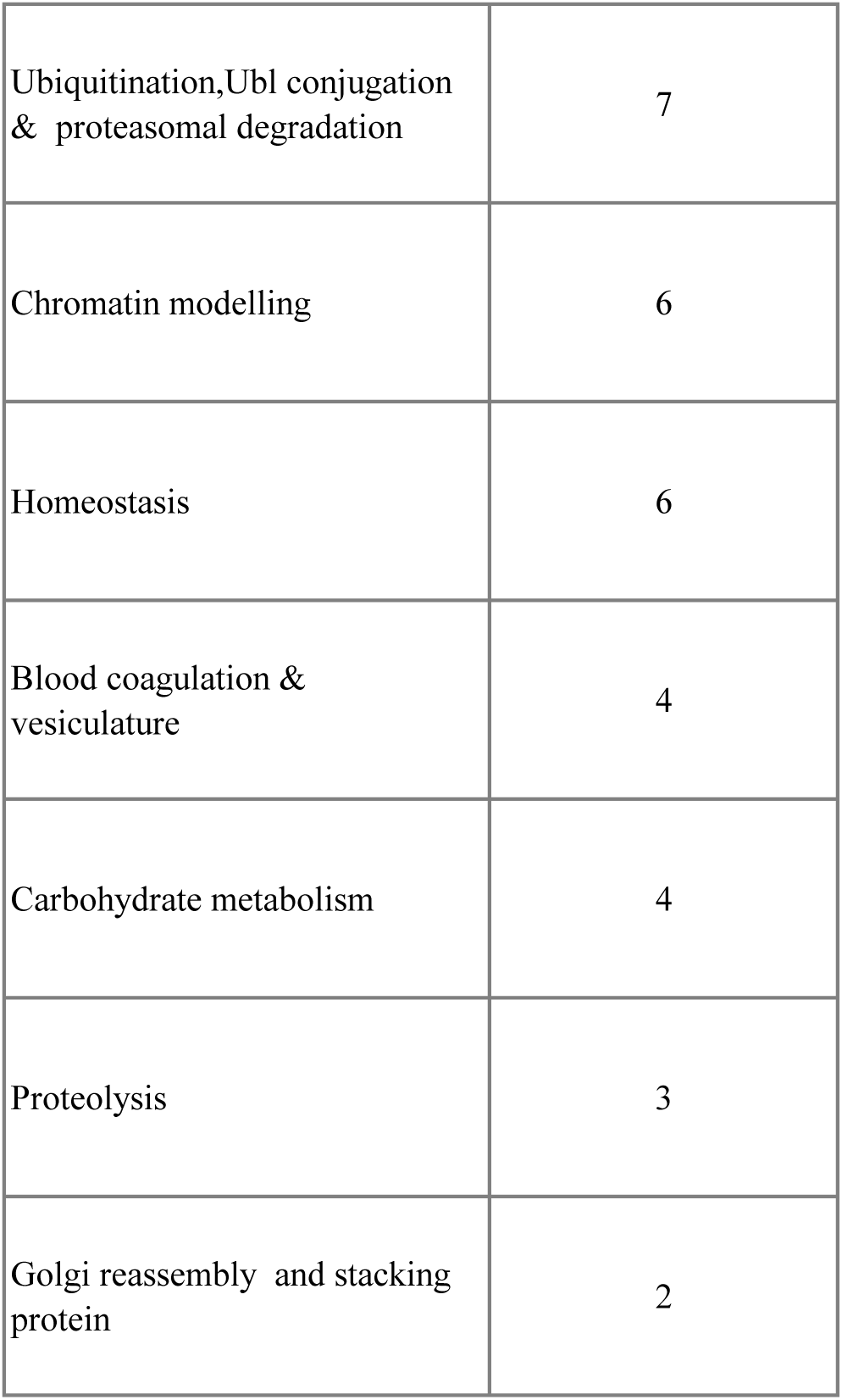
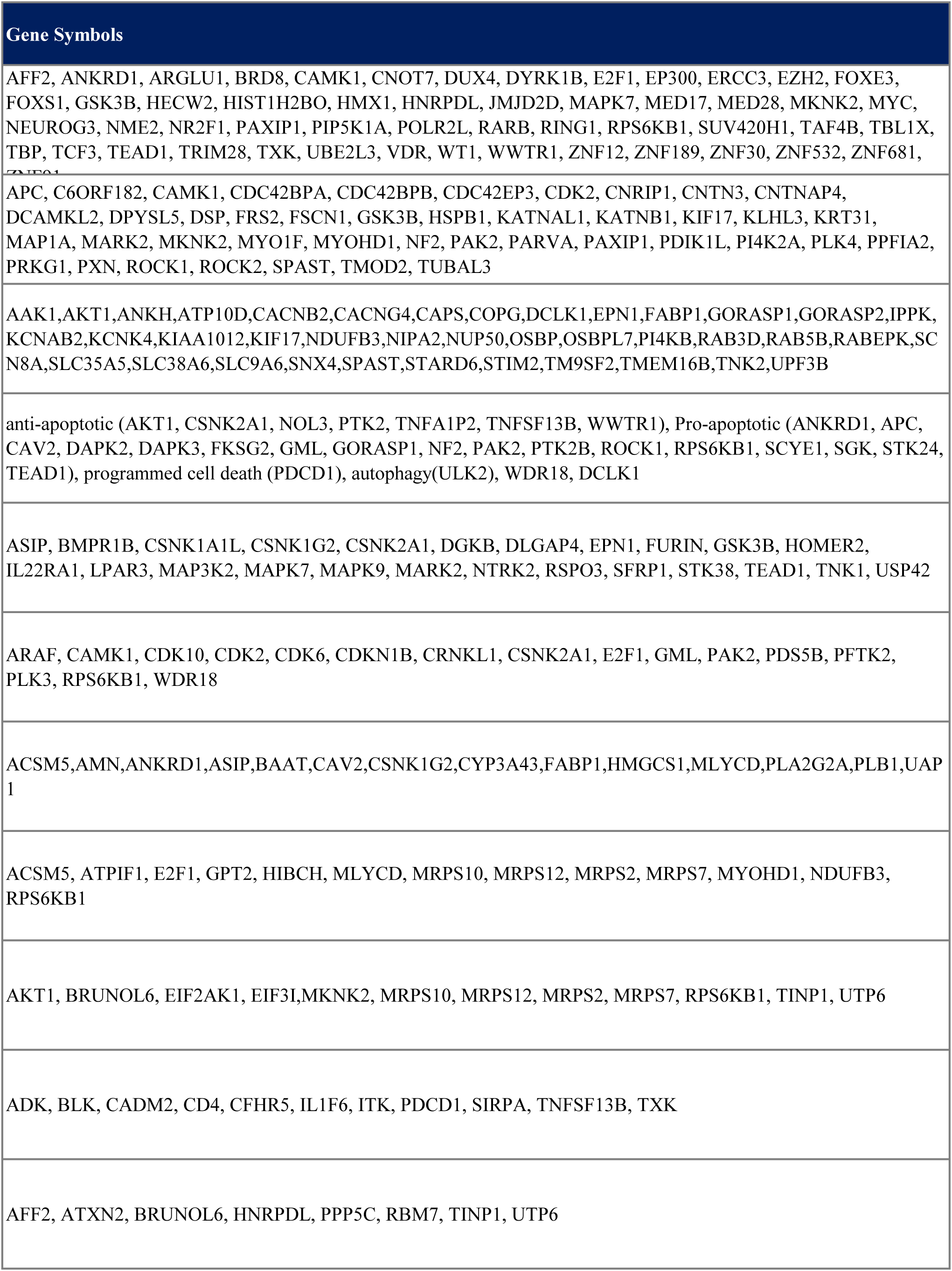
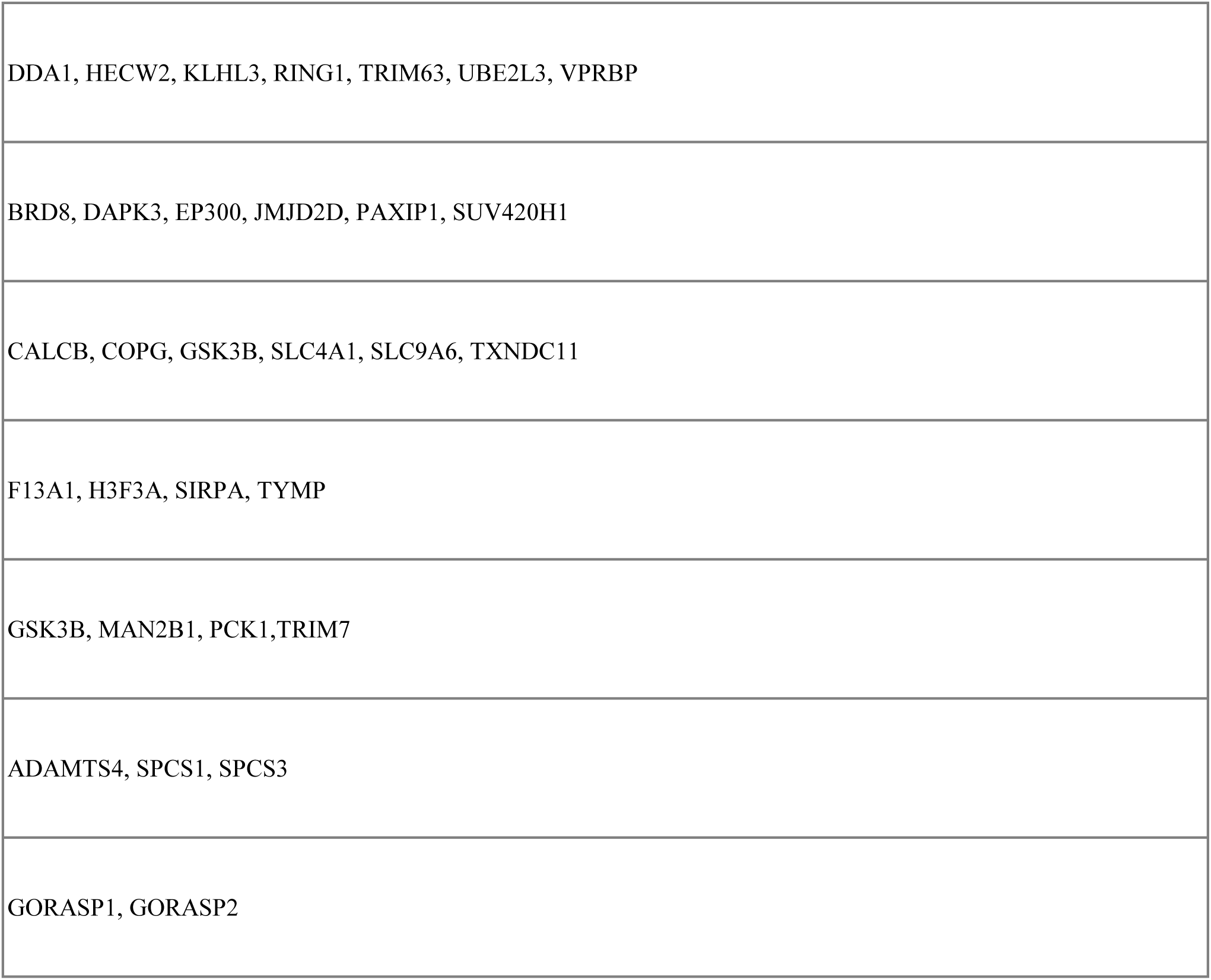
18 Functional Signatures and the inhibitors in each of the functional category are reported in this table.

